# Sub-minute phosphoregulation of cell-cycle systems during *Plasmodium* gamete formation

**DOI:** 10.1101/126276

**Authors:** Brandon M. Invergo, Mathieu Brochet, Lu Yu, Jyoti Choudhary, Pedro Beltrao, Oliver Billker

## Abstract

Malaria parasites are protists of the genus *Plasmodium*, whose transmission to mosquitoes is initiated by the production of gametes. Male gametogenesis is an extremely rapid process that is tightly controlled to produce eight flagellated microgametes from a single haploid gametocyte within 10 minutes after ingestion by a mosquito. Regulation of the cell cycle is poorly understood in divergent eukaryotes like *Plasmodium*, where the highly synchronous response of gametocytes to defined chemical and physical stimuli from the mosquito has proved to be a powerful model to identify specific phosphorylation events critical for cell-cycle progression. To reveal the wider network of phosphorylation signalling in a systematic and unbiased manner, we have measured a high-resolution time course of the phosphoproteome of *P. berghei* gametocytes during the first minute of gametogenesis. The data show an extremely broad response in which distinct cell-cycle events such as initiation of DNA replication and mitosis are rapidly induced and simultaneously regulated. We identify several protein kinases and phosphatases that are likely central in the gametogenesis signalling pathway and validate our analysis by investigating the phosphoproteomes of mutants in two of them, CDPK4 and SRPK1. We show these protein kinases to have distinct influences over the phosphorylation of similar downstream targets that are consistent with their distinct cellular functions, which is revealed by a detailed phenotypic analysis of an SRPK1 mutant. Together, the results show that key cell-cycle systems in *Plasmodium* undergo simultaneous and rapid phosphoregulation. We demonstrate how a highly resolved time-course of dynamic phosphorylation events can generate deep insights into the unusual cell biology of a divergent eukaryote, which serves as a model for an important group of human pathogens.

## Introduction

Malaria represents a major global health concern, causing an estimated 212 million cases and resulting in approximately 429 000 deaths in 2015 (World Health Organization, 2016). However, much of the fundamental biology of the causative agents, intracellular parasites of the eukaryotic genus *Plasmodium*, remains unknown. In particular, we lack fundamental knowledge about the key molecular systems that drive the parasites’ survival and transmission, due to significant evolutionary distance from well-studied model organisms and to experimental challenges posed by the parasite.

*Plasmodium* species have complex life cycles between a vertebrate host and a mosquito vector of the genus *Anopheles.* Ingestion of the developmentally arrested sexual-precursor stages, the gametocytes, by a susceptible mosquito initiates parasite transmission. Micro- and macrogametocytes respond to a small mosquito molecule, xanthurenic acid (XA), which must coincide with a drop in temperature to trigger their differentiation into male microgametes and female macrogametes (Billker et al., 1997). Cytosolic Ca^2+^ levels rise steeply after a lag phase of 6–8 s and peak within the first 20 s of activation (Billker et al., 2004). Within 15 s, a microtubule organising centre gives rise to eight kinetosomes in microgametocytes (Sinden et al., 1976). As soon as 60 s after activation, they have assembled monomeric tubulin from the nucleoplasm into the first mitotic spindle, and four axonemes start to grow on the templates of kinetosomes at each spindle pole (Billker et al., 2002). DNA replication in the microgametocyte is also initiated within the first 30 s after activation, *i.e.* before the first mitosis. The same is probably true for events leading to egress of both micro- and macrogametocytes from the host erythrocyte, since these are also dependent on the early Ca^2+^ signal (Billker et al., 2004). Within 8–10 min, each microgametocyte must replicate its genome three times, undergo three rounds of endomitosis, and assemble eight axonemes in order to produce eight microgametes. The microgametes then extrude from the gametocyte in a flagellar manner, through a process called exflagellation. Fertilisation of the macrogametes by microgametes results in the development of motile ookinetes, which may then escape the mosquito midgut.

The signal transduction pathway initiated through gametocyte activation by XA is an experimentally validated target for interventions to block parasite transmission to the mosquito (Ojo et al., 2012). A more detailed understanding of the mechanisms that link extracellular triggers to the cell- and developmental-cycle of the parasite is therefore of significant interest. While a parasite receptor for XA has remained elusive, a very early response of the gametocyte is thought to be a rise in intracellular cyclic guanosine monophosphate (cGMP) concentration through regulation of guanylyl cyclase and phosphodiesterase activity (Kawamoto et al., 1990; Muhia et al., 2001), causing the activation of the cGMP-dependent protein kinase PKG (McRobert et al., 2008). PKG activity stimulates the rapid release of intracellular Ca^2+^ stores via phosphoinositide metabolism (Brochet and Billker, 2016). This Ca^2+^ signal, in turn, directly regulates the protein kinase CDPK4 and the protein phosphatase calcineurin A (CnA), both of which are required for DNA replication during microgamete formation (Billker et al., 2004; Philip and Waters, 2015). CDPK4 is associated with the minichromosome maintenance (MCM) complex and its activity is required to initiate the assembly of the pre-replicative complex. Within the same period, CDPK4 activity is also needed to control the assembly of the first mitotic spindle (Fang et al., 2017). CDPK4 has separate, additional functions in cytokinesis and the activation of axoneme motility but it remains unknown whether these roles are initiated by a second Ca^2+^ signal (Fang et al., 2017).

While these data confirm the key role of CDPK4 in regulating the cell cycle during male gamete formation (Billker et al., 2004), other protein kinases and phosphatases are known to be involved. Another Ca^2+−^regulated kinase, CDPK1, may well be activated by the same Ca^2+^ signal but is only required for the egress of gametocytes of either sex from the host cell and in female gametes is also essential for the subsequent expression of translationally repressed mRNAs (Sebastian et al., 2012). A mitogen-activated protein kinase (MAPK2) has been implicated in multiple stages of gametogenesis, playing a role in mitosis, cytokinesis, and axoneme motility (Tewari et al., 2005; Rangarajan et al., 2005; Guttery et al., 2012). Two other enzymes, an SR-protein phosphorylating kinase SRPK1 (Tewari et al., 2010; Kern et al., 2014) and a protein phosphatase PPM1 (Guttery et al., 2014), have been found to be required to complete exflagellation, however more precise characterisations of their functions have not yet been made.

A comparison of phosphorylation events in a CDPK4 knockout mutant has identified a relatively small number of only 70 phosphosites that, within 18 s of gametocyte activation, differ significantly from wild type in their phosphorylation state (Fang et al., 2017). We hypothesised that a far greater number of differential regulation events can be revealed by a time-course experiment that would allow regulated sites to be differentiated according to the kinetics and direction of change. This would reveal a more comprehensive and unbiased network of co-regulated proteins in the different biological systems which are active during gametogenesis (Kanshin et al., 2015). To this aim we exploited the advantages offered by *P. berghei*, a parasite specific to rodent hosts, which allows gametocytes to be readily purified in sufficient quantities for biochemical investigation, and whose purified gametocytes can be activated rapidly and effectively *in vitro,* in a highly synchronous manner, through a drop in temperature and the concomitant addition of XA to the culture (Billker et al., 1997).

We have constructed a high-resolution phosphorylation time course of events happening during the first minute of gametocyte activation in *P. berghei*. Our results suggest that, in addition to CDPK4, other lipid and protein kinases contribute to the early gametogenesis signalling response. We show that systems that are fundamental to microgamete formation, particularly the assembly of axonemes, the formation of the mitotic spindle and DNA replication-initiation and replication, undergo phosphoregulation almost instantaneously and simultaneously during the first seconds of cell activation. We validate our analysis using the phosphoproteomes of both CDPK4 and SRPK1 mutants and further identify a new role for the latter in gametogenesis. We also present a resource of hundreds of regulated phosphosites of likely importance for controlling basic biological processes in malaria parasites.

## Results

### Robust quantification of phosphosites

Activation of gametocytes by XA at a permissive temperature is very rapid (Figure 1a). We therefore focused our time course on the first 18 s, at 6 s intervals. We expected this to capture the first cellular response to the initial Ca^2+^ signal. The time course finishes after the first minute, since this period should be sufficient to capture the key regulatory events associated with the induction of major cell cycle events in the microgametocyte.

For each time point, we used four label-free biological replicates and produced two technical replicates from each phosphopeptide-enriched sample (Figure 1b). A total of 17 238 unique peptides were identified, of which 8982 were phosphopeptides. 12 672 phosphosites were identified within these peptides, with 10 237 localised at site-level with high-confidence (confidence > 0.75); 8299 of these high-confidence sites were found on *P. berghei* proteins and 1938 were from *M. musculus* proteins (Supplementary Table S1). Phosphosite intensities, estimated from the intensities of the different phosphopeptides on which the sites are identified, were strongly correlated between replicates (Figure 1c). The data covers several of the proteins and phosphosites observed in a previous, two-dimensional gel electrophoretic screen of phosphorylation in gametocytes and gametes (Alonso-Morales et al., 2015). Thus, the data represents a comprehensive and reproducible view of early phosphorylation during gametogenesis, from which we could reliably reconstruct time courses.

**Figure 1:**
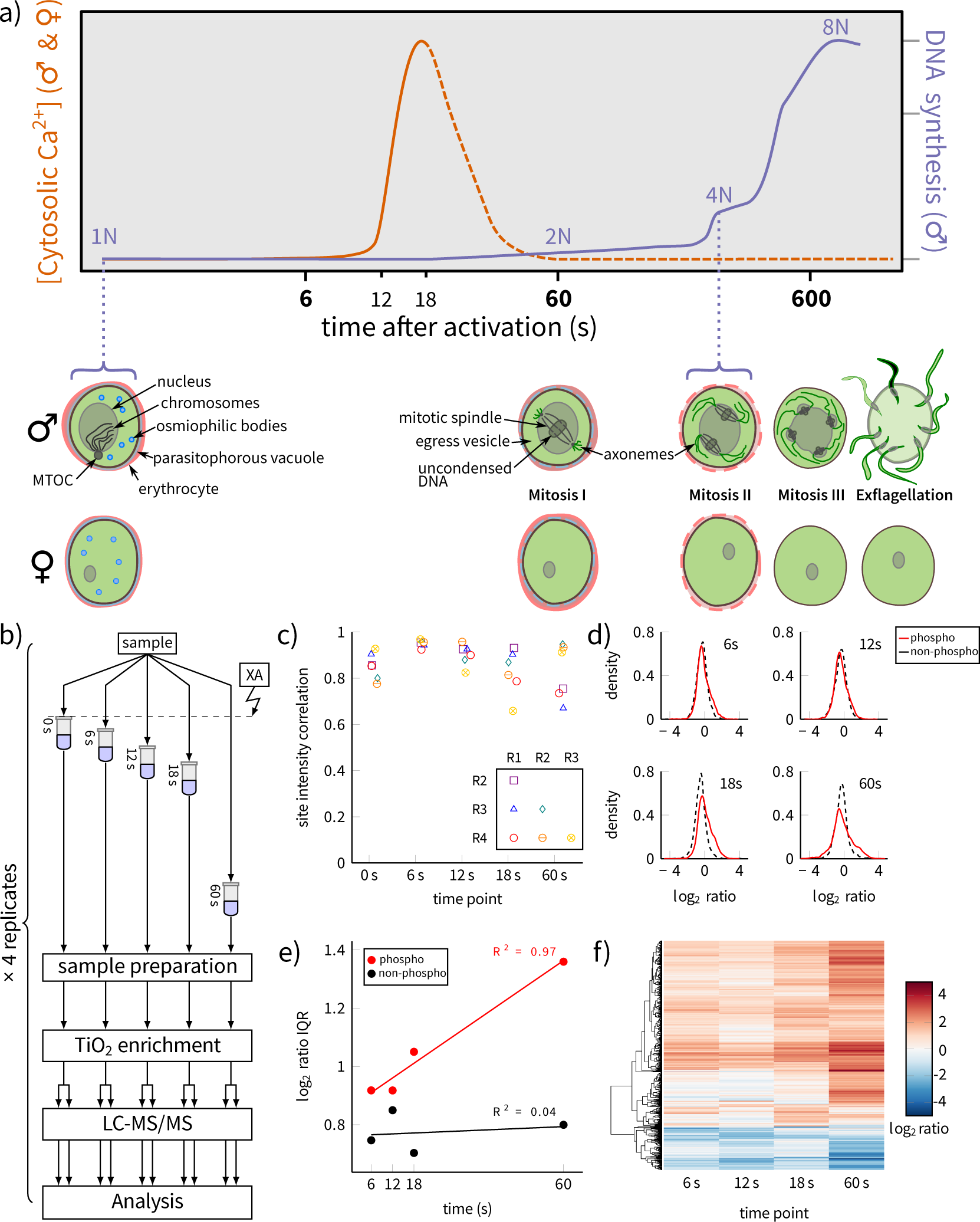
a) Timing of key events during *P. berghei* gametogenesis based on published data. Rapid mobilisation within 20 s of intracellular Ca^2+^ stores leads to egress from the parasitophorous vacuole and the erythrocyte in both micro- (♂) and macrogametocytes (♀). Microgametocytes also produce eight axonemes and undergo three rounds of DNA replication and endomitosis within 10 min, followed by exflagellation. (MTOC: microtubule organising centre; Ca^2+^ and DNA synthesis dynamics were adapted from Billker et al. (2004) and Raabe et al. (2009), respectively) b) A schematic of the experimental design. c) Correlation coefficients for phosphosite intensities between biological replicates for each time point (R1: replicate 1, R2: replicate 2, *etc.*). d) Kernel density distributions of log_2_-ratios for phosphosites and non-phosphopeptides at the 6, 12, 18 and 60 s time points relative to unstimulated gametocytes (0 s). The non-phosphopeptide ratio distributions were used as null-distributions to test significance of the phosphosite ratios. e) The interquartile ranges (IQRs) of log_2_-ratio distributions from (d), with least-square regression lines and Pearson correlation coefficients shown. The increase in IQR for phosphosites represents the phosphorylation signal spreading out to a wider range of targets over time. f) Hierarchical clustering of log_2_-ratios of the significantly regulated phosphosites.

## Phosphorylation time courses cluster into distinct response groups

We quantified the change in phosphorylation states over time by calculating log_2_-ratios against unstimulated gametocytes (0 s), producing time courses of changes in abundance from 0–60 s. In order to account for sites which were not detectable as phosphorylated at the beginning or at the end of the experiment, we also constructed two “truncated” time courses: 6–60 s time courses for sites that were not observed at 0 s and 0–18 s time courses for those that were not observed at 60 s.

1089 phosphosites on 549 *P. berghei* proteins were found to show evidence of significant change compared to non-phosphopeptides (Figure 1d) during one of the three time courses (referred to herein as “regulated sites”, however it is important to emphasise that this is in reference to regulation of phosphorylation state and may not in all cases correspond to regulation of protein activity). The distributions of fold-changes were found to widen over time, meaning that more sites were significantly regulated as the activation process progressed (Figure 1e). This has been interpreted as the signalling response spreading out to a wider range of targets over the course of the experiment (Kanshin et al., 2015). For most regulated sites, full, 0–60 s time courses were measurable (0–60 s: 926 sites; 6–60 s: 64 sites; 0–18 s: 99 sites; Supplementary Table S2). Hierarchical clustering of the time courses revealed general up-regulation trends for the majority of the sites and down-regulation for only about a fifth of the sites (Figure 1f).

To gain finer detail, we employed a Gaussian mixture-model clustering algorithm to re-partition the data (Figure 2a; see also Supplementary Table S2). Using this method, the 0–60 s time courses were partitioned into eight clusters: seven clusters that show general up-regulation and one showing down-regulation. Clusters 1, 2, and 3 consist of sites that are already up-regulated within 6–12 s, while clusters 4, 5, and 6 show predominantly later up-regulation. Cluster 7 contains up-regulated sites that did not fit well in the other clusters. For the truncated time courses, the algorithm produced two clusters each (one up-regulation and one down-regulation cluster) (Figure 2a; 6–18 s: clusters 8 and 11; 0–18 s: clusters 9 and 12). Through this clustering of the time course results, we see that the phosphosites in the parasite are characterised by several classes of dynamic responses to gametocyte activation, with many displaying activity within the first seconds after the initial stimulus.

**Figure 2:**
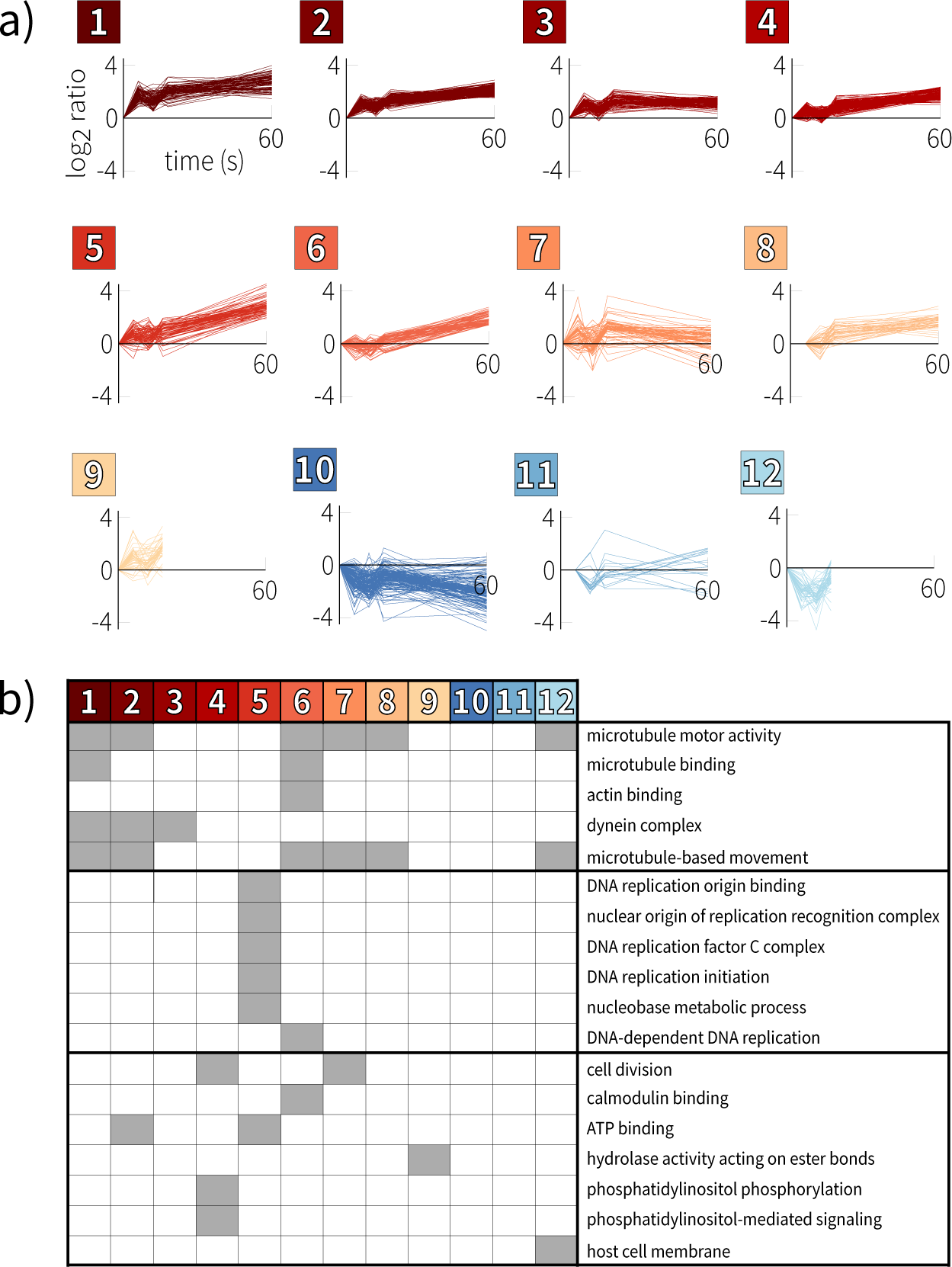
a) Gaussian mixture-model-based clustering of *P. berghei* phosphorylation time courses. b) Enriched Gene Ontology terms for the *P. berghei* time-course clusters.

## DNA-replication and microtubule-related proteins are primary targets of phosphoregulation

Of the 549 significantly regulated proteins, 436 were annotated with at least one GO term, which we used to detect biological functions enriched within individual clusters (Figure 2b; see Supplementary Figure 1 for enrichment in higher-level parent terms and Supplementary Table S3 for test statistics); all significant terms have *p* < 0.05 after correcting for multiple testing). This analysis was entirely consistent with the known importance of cell-cycle events during male gametogenesis. Microtubule-related terms were associated with clusters of different phosphorylation kinetics during the first minute after gametocyte activation (compare for instance clusters 1 and 6 in Figure 2a, which are both enriched in proteins annotated for microtubule motor activity), potentially reflecting different roles in spindle and axoneme assembly, while proteins functioning in DNA replication-initiation and nucleic acid metabolism were only moderately phosphorylated within the first 18 s (cluster 5), and their phosphorylation continued to increase after 18 s (clusters 5 and 6). One early and one late up-regulation cluster (clusters 2 and 6) were also enriched in male-specific proteins (*p* = 0.035 and *p* = 0.023 respectively), while another early up-regulation cluster (cluster 3) was enriched for female-specific proteins (*p* < 0.001) and the primary down-regulation cluster (cluster 10) was enriched for proteins that are exclusive to neither male nor female gametocytes (*p* = 0.035).

Our global phosphorylation analysis was in good agreement with our recent study focused on CDPK4 (Fang et al., 2017). Microgametocytes enter S-phase I within 30 s of activation and, accordingly, CDPK4 and two of its known substrates carry sites that are up-regulated early (cluster 3). In both cases, this included the exact residues targeted directly by CDPK4, *i.e.* S9 in PBANKA_072020 (SOC1) and S5051 in PBANKA_144220 (SOC2). Interestingly, the biological functions associated with these substrates are quite different, with a knock out of SOC1 indicating a role in initiation of replication, while SOC2 was crucial to assemble the mitotic spindle during microgametogenesis (Fang et al., 2017).

Within the first minute of microgametogenesis, spindle plaques need to split, nucleate the spindle of mitosis I, give rise to basal bodies and start growing axonemes. Regulation of mitotic spindle assembly remains poorly characterised in *Plasmodium.* We find SOC2 has additional phosphosites upregulated later (clusters 6 and 7). SOC2 may thus integrate multiple signals from different kinases to regulate the stability of the mitotic spindle. In other eukaryotes, microtubules are nucleated from a ring of γ-tubulin in complex with a family of interacting proteins, many of which are phosphorylated by mitotic protein kinases (Teixidó-Travesa et al., 2012). In *Plasmodium*, only γ-tubulin and one component of the ring complex, PBANKA_083430, appear to be conserved. The latter is characterised in our data by an early phosphorylation event from 6 s (cluster 5; albeit showing an atypical peak at 6 s followed by a gradual recovery by 60 s). Similarly, phosphosites in the centrosome component centrin-2, in a number of axonemal dynein chains and in other axoneme associated proteins (Talman et al., 2014) are also contained in the rapid-response clusters 1–3.

Interestingly, while phosphatidylinositol (PI) signalling is known to be triggered early by PKG and is required for the rapid mobilisation of Ca^2+^ (Brochet et al., 2014), we saw enrichment for PI metabolism in sites which show predominantly late up-regulation (cluster 4). Closer inspection of the data reveals a more differentiated picture, however. This cluster, indeed, includes phosphosites on a PI 3-kinase (PI3K, PBANKA_111490), a PI 4-phosphate-5-kinase (PI4P5K, PBANKA_020310) and a PI 4-kinase (PI4K, PBANKA_110940). The PI4K, a validated drug target (McNamara et al., 2013), carries two regulated sites that are both up-regulated within 6 s, as would be predicted: S534 (cluster 2) and S538 (the aforementioned site in cluster 4). However, S538 has more complex dynamics, with its initial phosphorylation suppressed by 18 s and then reappearing by 60 s, hence its inclusion in the late up-regulation cluster. Later phosphorylation events in the pathway such as this may be involved in signal termination or restoration of IP3 levels. It is noteworthy that in another life-cycle stage of *P. berghei*, the motile ookinete, the same sites were identified as direct or indirect targets for PKG and were shown to be of functional importance during ookinete gliding (Brochet et al., 2014).

Taken together, these results indicate that our dataset covers most of the few phosphorylation events known to be involved in the regulation of the *Plasmodium* cell cycle and should thus contain other functionally relevant events. The functional enrichment analysis highlights high-level trends that are congruent with our limited existing knowledge of gametogenesis and further indicates that mitotic proteins may become active at the same time that DNA replication is initiated, not after the cell has completed S-phase.

## The phosphoregulated proteins are functionally associated

We next investigated whether the regulated phosphoproteins tend to be functionally related to each other in a previously published protein-protein association network (Szklarczyk et al., 2015). Functional associations are predicted from diverse data such as co-expression or co-evolutionary patterns. We found that the subset of the network consisting of phosphoregulated proteins is more highly interconnected than expected by random chance (Supplementary Figure 2). We then visualised the network of regulated proteins that were associated with the enriched functional terms described in the previous section (Figure 3). If the phosphoregulated proteins represented a diverse set of systems, then this network would be composed of several disconnected or loosely connected components. However, of the 124 proteins present in the network and annotated with the functional terms, the largest connected component of the network included 102 of them. In other words, most of the proteins that are phosphoregulated during gametogenesis are functionally associated with each other. The network consists of two major clusters of proteins (Figure 3). One is composed of proteins annotated for microtubule motor activity and which tend to be phosphoregulated at many different sites. About a third of these motor proteins have previously been implicated as components of the axoneme (Talman et al., 2014), while the rest are likely to be related to the mitotic spindle. The other major cluster consists largely of helicases, protein kinases, and DNA-binding proteins, comprising the replisome. Other minor clusters are present, including one of ribosomal proteins and one related to phosphoinositide signalling. These clusters are heavily interconnected via a CDK-related (cyclin-dependent kinase) protein kinase, CRK5 (PBANKA_123020). An orphan kinase PBANKA_062210, found within the replisome cluster, also holds a very central position among these proteins. This tight connection between phosphoregulated proteins in a replisome cluster and a microtubule cytoskeleton cluster suggests that regulation of the cell cycle following microgametocyte activation is a result of unified and coordinated signalling events.

## Protein kinases and phosphatases associate with specific dynamic clusters

Given the presence of phosphoregulated kinases in our data, we next asked whether these kinases are responsible for the general phosphorylation patterns that we observe. Because the cell cycle in the *Plasmodium* sexual stages is likely to be orchestrated by both stage-specific and persistent regulators, many of which are intractable by reverse genetics, we could potentially discover otherwise difficult-to-obtain functional information through the time-course data. By using, on the one hand, the protein-protein association network to infer functional relationships between proteins and, on the other hand, our phosphosite clusters to identify groups of proteins undergoing phosphoregulation with similar dynamics, we can statistically test whether the kinases are significantly associated with any of these regulated protein groups (see Methods). By furthermore considering the functional enrichment of the phosphosite clusters, we can construct a coarse-grained view of putative functional roles of the phospho-signalling enzymes.

An aurora kinase, ARK2 (PBANKA_040740), was observed to have two up-regulated phosphosites. Accordingly, it associates on the network with a significant number of proteins that had either rapidly phosphorylated sites (cluster 1) or late-phosphorylated sites (cluster 6) (Figure 4, top-left). These two phosphosite clusters were enriched in microtubule motor-proteins (Figure 2b), many of which are functionally associated with ARK2, with little overlap between the two sets of proteins. This links ARK2 to the phosphoregulation of the microtubule motor proteins, which is consistent with another member of the aurora kinase family being found at microtubule organising centres in *P. falciparum* (Carvalho et al., 2013). The late up-regulation cluster (cluster 6) was also enriched in male-specific proteins, thus, we hypothesise that ARK2 plays a role in regulating mitotic processes or axoneme formation during microgametogenesis, possibly integrating different signals over the course of the process.

**Figure 3:**
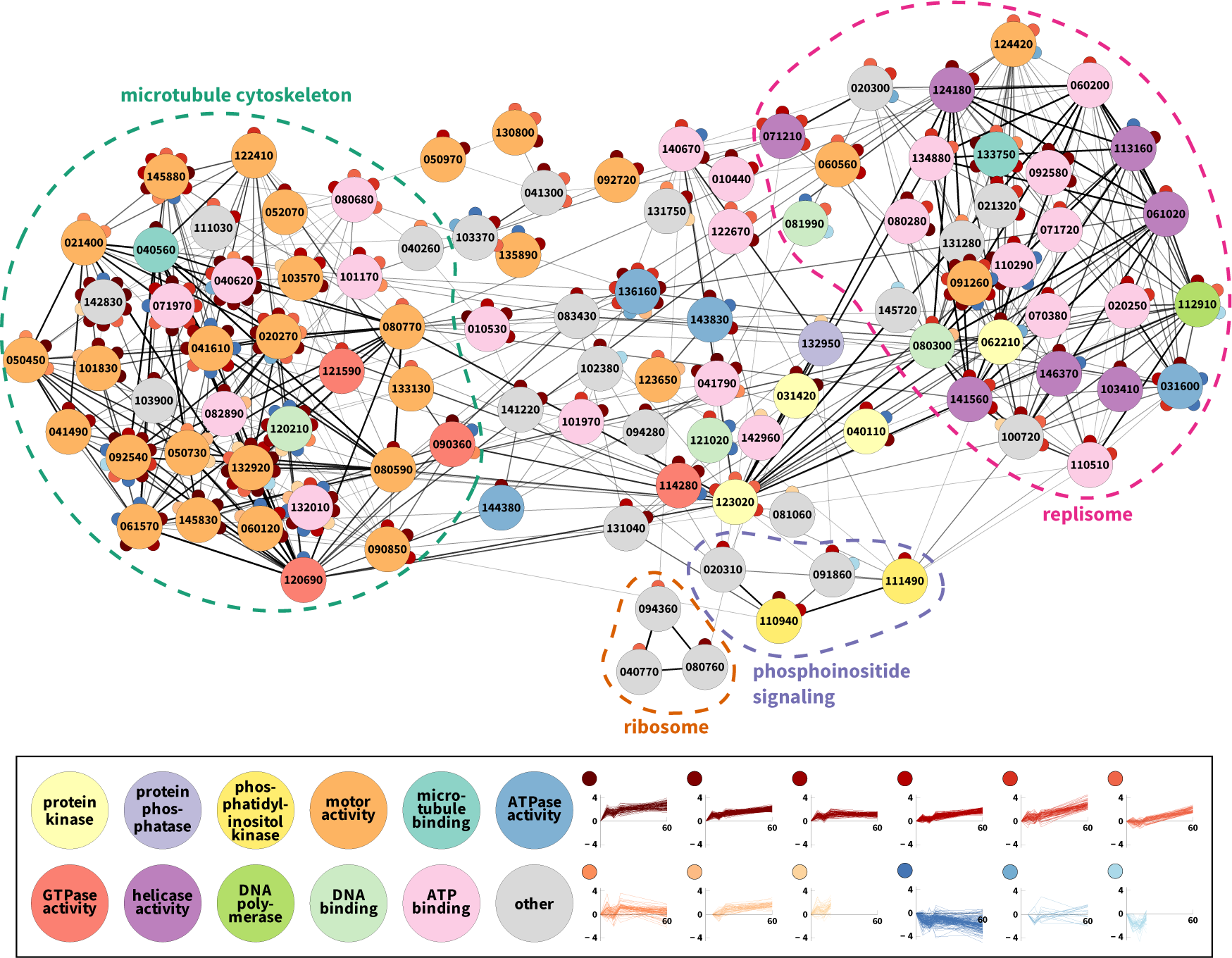
The proteins that are phosphoregulated following gametocyte activation have evidence of tightly interconnected functional associations. The subset of the functional-association network consisting of regulated phosphoproteins that were associated with enriched GO terms (minimum edge score of 300). Edges are shaded and weighted according to their edge score. Nodes are coloured according to molecular function annotations and contain the numeric portion of the protein’s ID. Sub-nodes represent significantly regulated phosphosites, with their colour reflecting their time course cluster.

The kinase NEK1 (PBANKA_144300), thought to be involved in MAPK signalling, is known to be essential and is expressed highly in males at the gametocyte stage (Khan et al., 2005; Tewari et al., 2010; Dorin-Semblat et al., 2011; Carvalho et al., 2013; Tao et al., 2014). Following gametocye activation, it was found to be phosphoregulated at one site, characterised by rapid dephosphorylation within 6 s followed by recovery to pre-activation phosphorylation levels by 18 s. On the network, it shows a pattern of association with proteins whose phosphoregulation is similar to that of the kinase (cluster 11) (Figure 4, middle-left). Thus, we would speculate that NEK1 is primarily active by 60 s and indeed may be functionally repressed during early signalling events. This is consistent with the role of MAPK signalling during later gametogenesis events (Tewari et al., 2005).

In a more complicated case, we found overlapping patterns of functional associations between CRK5 (PBANKA_123020), the orphan kinase PBANKA_062210, and a putative protein phosphatase 2C (PBANKA_142720). CRK5 was up-regulated at multiple sites following gametocyte activation, while PBANKA_062210 and the phosphatase had similar patterns of up- and down-regulation at two sites. All three enzymes are associated with proteins carrying phosphosites that are predominantly up-regulated at later time-points, with some early activity (cluster 5) (Figure 4, bottom-left). This links the enzymes with DNA replication initiation (Figure 2b). Fittingly, in our network-clustering analysis, CRK5 and PBANKA_062210 were found to be tightly associated with the replisome, highlighting the potential important of these kinases (Figure 3; the phosphatase is not represented in the figure). CRK5 and the phosphatase also have significant interactions with proteins undergoing broad dephosphorylation (cluster 10). Lastly, the phosphatase is associated with proteins that undergo late up-regulation (cluster 8), which is enriched in microtubule motor proteins (Figure 2b). It is also interesting to note that CRK5 is among the late-up-regulated proteins that are functionally associated with ARK2, suggesting that the regulatory functionality of ARK2 and CRK5 may, too, be related. Such inter-phospho-enzyme associations further highlight the probability that we have observed a single, broadly functioning signalling module that coordinates cell-cycle progression in gametocytes.

This analysis places these phospho-enzymes alongside others that have been more directly implicated in gametogenesis through knock-out or loss-of-function experiments (Figure 4). However, beyond the initial events around the second messengers cGMP and Ca^2+^, we cannot yet derive a hierarchical structure between the enzymes in order to reconstruct a gametogenesis signalling pathway. The second-messenger dynamics suggest that the signal originates with PKG and the Ca^2+^-dependent enzymes, but specific substrates of these and other enzymes remain largely unknown. In *P. falciparum*, CDPK1 has been identified as a substrate of PKG, so we indicate this relationship in Figure 4 (Alam et al., 2015). Similarly, the NIMA-related kinases NEK1 and NEK3 of *P. falciparum* have been shown to be able to phosphorylate MAPK2 *in vitro*, suggesting that they could regulate the functionality of this kinase (Lye et al., 2006; Dorin-Semblat et al., 2011). This, in turn, would tentatively link one phosphosite cluster (cluster 11) to mitosis and axoneme motility via NEK1 and MAPK2 (Figure 4). Surprisingly, none of the previously implicated phospho-enzymes were otherwise found to be statistically associated with the time-course clusters. This is likely due to low coverage of the network, in which many of the enzymes, such as CDPK4 and SRPK1, have relatively few high-confidence interactions. Nevertheless, the results introduce several new enzymes as likely components of the pathway, with their specific roles requiring further validation.

## Conservation of phosphosites suggests functional constraint

If a phosphosite has a vital biological function, it is likely to be conserved at the sequence level and phosphorylation at homologous sites should be observable in different species. We assessed the sequence conservation of phosphosites at the underlying nucleotide level (dN/dS; see Methods) per site during *Plasmodium* evolution (Supplementary Table S4). We defined “conserved sites” as those that show evidence of more selective constraint (lower dN/dS) than would be predicted by known determinants of evolutionary rates. The proteins with conserved phosphosites were enriched for motor activity, the myosin complex, calmodulin binding, and cytoskeletal protein binding. When limited to only proteins undergoing active phosphoregulation during gametogenesis, the set is enriched in proteins annotated for DNA helicase activity, DNA replication and replication-initiation, and the MCM complex.

We then focused on the conserved, regulated sites that fall within a predicted protein domain and which have previously been detected as phosphorylated in *P. falciparum*. Together, these would be likely indicators of important functionality. We identified thirteen such sites (Table 1). Notably, both CDPK1 and CDPK4 each have one such phosphosite located within their kinase domains, suggesting that these sites can be taken as indicators of the respective kinase’s activity. This is supported by the fact that the orthologous CDPK1 site in *P. falciparum* has been identified as a site of autophosphorylation (Ahmed et al., 2012). Several sites also fall within domains related to DNA replication and the cell cycle, for which evidence in other species suggests some functionality. In particular, phosphorylation within the MCM N-terminal domain of human MCM7 has been implicated in the regulation of formation of the MCM2-7 complex and in the progression of the cell cycle (Wei et al., 2013) and phosphorylation within the MCM domain of human MCM3 regulates the replication process (Han et al., 2015). The phosphorylation of MAT1 (PBANKA_111190) within its eponymous domain is also potentially related to the regulation of the cell cycle: in other eukaryotes, this protein (MNAT1 in humans) associates with CDK7 and cyclin H to form the CDK-activating kinase enzymatic complex (CAK), which then regulates cyclin-dependent kinases by phosphorylating them. However, no functionality of phosphoregulation of MNAT1 has yet been described.

**Figure 4:**
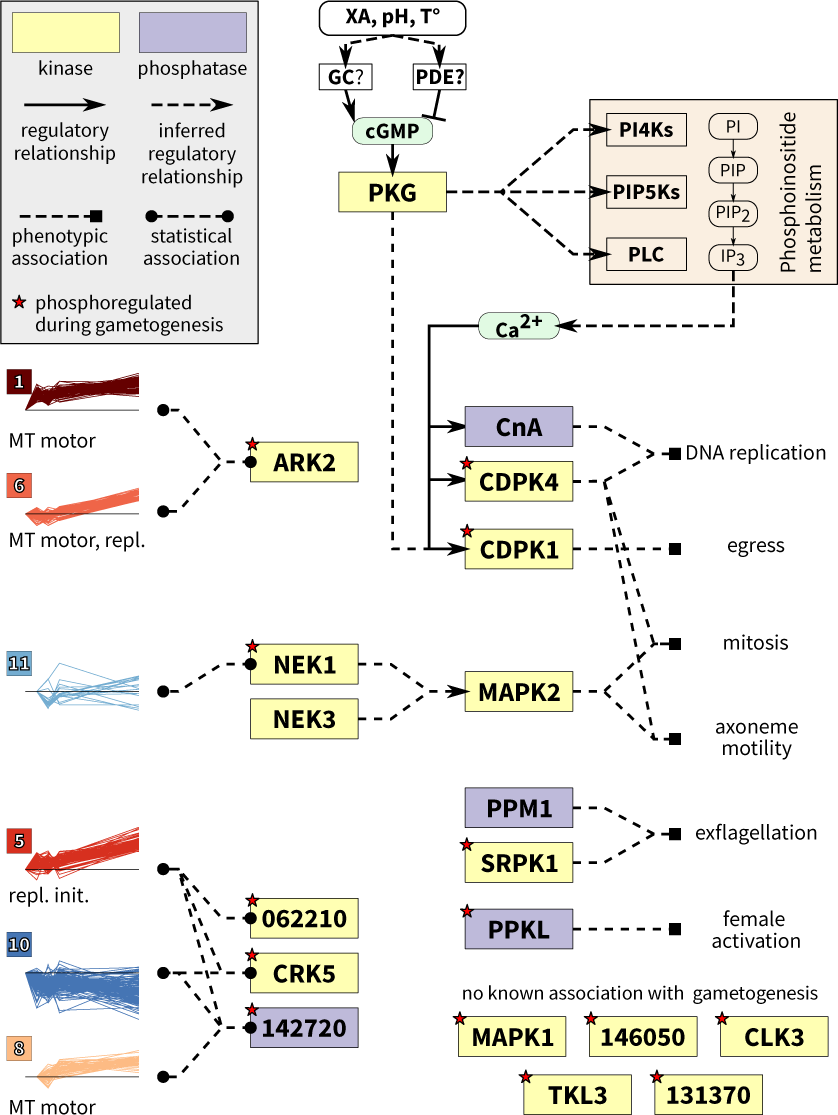
A schematic of early phosphorylation signalling during gamete formation. Several of the phosphoregulated kinases and phosphatases can be associated with the phosphosite clusters via statistical enrichment on a protein functional-association network; functional terms enriched in each cluster are listed (MT motor: microtubule motor activity; repl. init.: replication initiation; repl.: DNA replication). Other enzymes have been associated with specific stages of gametogenesis through knock-out or loss-of-function experiments. Inferred regulatory relationships include cases in which a kinase is known to be able to phosphorylate another kinase, but no specific function has been described for the event.

**Table 1:**
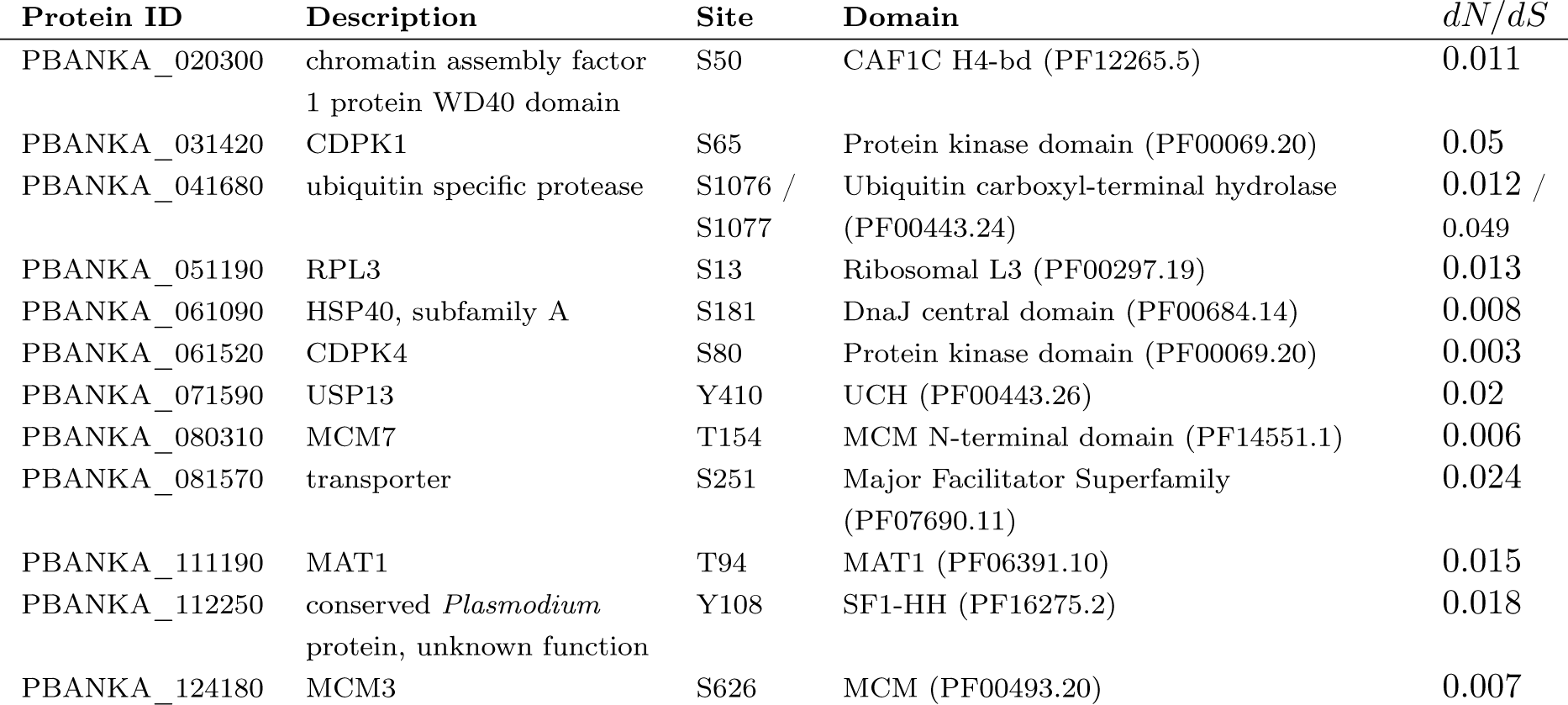
Significantly regulated sites which show stronger conservation, lie within a predicted protein domain, and have been previously observed in *P. falciparum,* which we take to be indicators of likely functional importance.

We were also intrigued by the large number of phosphoregulated sites observed on individual motor proteins. To determine whether these patterns are significant, we checked if the sites fall in so-called “phosphorylation hot-spots,” homologous regions within a protein family that are frequently found to be phosphorylated (Beltrao et al., 2012). If the phosphosites regulated during gametogenesis occur within such a hot-spot, then it may be an indication of functional importance. In a dynein light intermediate chain (PBANKA_041490), one of the two regulated phosphosites, T629 (Supplementary Figure 3a alignment position 850), falls in a major C-terminal phosphosite hot-spot. The orthologs in humans (Uniprot ID O43237) and mice (Q6PDL0) have phosphoserines aligned with this position (S383 in both species), however no function for this site is known. On the other hand, a N-terminal phosphorylation hot-spot in the dynein intermediate chain family has a known regulatory role (Vaughan et al., 2001; Towns et al., 2009; Bader et al., 2011; Rosse et al., 2012; Blasier et al., 2014; Young et al., 2015). One site near this region on PBANKA_050450 was found to undergo regulation during gametogenesis (S120; Supplementary Figure 3b alignment position 232). In human cells, phosphorylation of intermediate chain 1 within this hot-spot was found to mediate cargo binding in organelle transport(Vaughan et al., 2001; Towns et al., 2009). The dynein heavy chain and kinesin families were found both to be heavily phosphorylated along their full lengths, however no hot-spots were distinguishable in the dynein heavy chain family (Supplementary Figure 3c & d). In the kinesin family, several regulated sites fall within hot-spots of unknown function at well-aligned positions, particularly near the N- and C-termini. Thus, while we do not have functional knowledge of all motor-protein hot-spots, the existence of conserved phosphorylation patterns across diverse species points to an important role of phosphorylation in regulating these proteins’ functionalities.

## Gene knock-outs link protein kinases to core signalling responses

We next investigated the roles of specific protein kinases in the gametocyte activation response. We studied two knock-out lines, one lacking CDPK4 and the other with a deletion of SRPK1. Both kinase mutants make morphologically normal gametocytes but are defective in male gametogenesis (Billker et al., 2004; Tewari et al., 2010). While CDPK4 has been the subject of recent targeted analysis (Fang et al., 2017), much less is known about the specific activities of SRPK1. Consistent with these genetically confirmed functions, both undergo phospho-regulation during our time course. However, we could not derive any new functional insight into the kinases via statistical analyses, due to their poor connectivity on the network. We thus employed the knock-out lines to compare phosphorylation patterns before and 18 s after activation, with and without each kinase. We confirmed the reliability of the results by comparing to known functionality for CDPK4 and then focused on new observations in SRPK1.

In the CDPK4 knock-out experiment (CDPK4-KO), we quantified 3539 unique peptides in the wild-type (WT) and knock-out (KO) samples, belonging to 993 *P. berghei* proteins and 663 *M. musculus* proteins (Supplementary Table S5). Of the phosphosites found to be under significant regulation in the time-course experiment, 190 were unambiguously quantified in CDPK4-KO. In the SRPK1 knock-out experiment (SRPK1-KO), 3949 peptides were quantified from 1060 *P. berghei* proteins and 787 host proteins (Supplementary Table S5). 236 peptides carried sites that were observed as regulated in the time-course experiment. The small overlap between the different types of experiments can be accounted for by the inherent stochasticity of MS peptide detection in complex biological samples and potentially by the difference in time points measured.

We saw significant differences between KO and WT after activation in both experiments, however in SRPK1-KO we also saw differences between the two samples before activation (Figure 5a). To investigate this, we first checked that the WT samples replicated the observations of the time course experiment. The significantly up- and down-regulated WT phosphopeptides in both experiments were enriched in protein functions that generally matched those of the time course experiment, despite the shorter time-frame (Figure 5b). We therefore concluded that the WT samples reproduced the observations from the time course experiment.

Pre-activation differences between KO and WT may have been caused by accumulated perturbations to protein abundances or phosphorylation during gametocyte formation, which would persist during gametogenesis if not under further regulation by the kinase. Indeed, for both experiments, we found a positive correlation between pre- and post-activation disruptions (Figure 5a). We interpreted outliers to this correlation as being specifically dependent on the kinase during gametocyte activation (Figure 5a, orange points). In CDPK4-KO, we found 196 dependent phosphosites, most of which were negatively disrupted in the KO. In SRPK1-KO, we identified 243 dependent phosphosites, the majority of which were positively disrupted. 16 of the SRPK1-KO disrupted sites were observed in the time course, assigned to both up- and down-regulation clusters, while 22 CDPK4-dependent sites were observed in the time course, all from up-regulation clusters. Thus, deleting CDPK4 leads to a significant loss of phosphorylation events, while a lack of SRPK1 produces notable gains thereof as well as a loss of dephosphorylation events. This suggests SRPK1 may indirectly repress phosphorylation via an intermediary enzyme.

We next performed a functional enrichment analysis on the disrupted proteins (Figure 5b). We found an enrichment in proteins related to the microtubule cytoskeleton in negatively disrupted proteins in CDPK4-KO and in both positively and negatively disrupted proteins in SRPK1-KO. Specifically, in the case of CDPK4 the phosphosites were up-regulated in WT and unregulated in the KO after activation; while for SRPK1 the trend was reversed (Figure 5c). Notably, the specific phosphosites that were disrupted were unique in each experiment. Thus, CDPK4 and SRPK1 appear to both regulate different subsets of the microtubule motor system and the phosphorylation patterns support the hypothesis of a repressive function for SRPK1. We also saw a disruption in both experiments of DNA replication origin binding, through components of the origin of replication initiation complex (ORC) and the MCM complex (Figure 5b). One interesting case is S60 on ORC1 (PBANKA_060200), which is up-regulated after activation in WT. When CDPK4 is not present, regulation of this site is lost. However, when SRPK1 is not present, the site significantly increases in abundance relative to WT. This suggests that the signal for phosphorylation of this site originates with CDPK4 and the stoichiometry of this event is moderated by the signal passing through an SRPK1-mediated feedback loop. This is supported by the observation that phosphorylation of SRPK1 itself is negatively disrupted when CDPK4 is deleted, placing SRPK1 downstream of CDPK4 in the early gametogenesis signalling events.

The phosphoproteomic effects of deleting CDPK4 fit our general understanding of the role of this kinase following gametocyte activation and thus impart confidence in the SRPK1-KO results. In order to place the SRPK1-KO phosphoproteomic results in a broader phenotypic context, we measured DNA replication, axoneme assembly and exflagellation patterns of the mutant parasites. We found that, by 10 min, a significantly lower percentage of gametocytes completed three rounds of DNA replication, while a larger percentage remained haploid (Figure 5d). This indicates that SRPK1-KO male gametocytes are affected in the first genome replication, but once this latter happens, no further defects are observed in reaching the octoploid level. Consistently, we observed that parasites that reached the octoploid level successfully assembled axonemes (Figure 5d and e). While neither observation could be accounted for by a change in sex-ratio in the mutant parasites (Figure 5d), they may be due to some proportion of the microgametocyte population becoming nonviable during gametocytogenesis. This is tentatively supported by our SRPK1-KO phosphoproteomic results, which indicated strong pre-activation disruption. However, the data also revealed significant post-activation effects on phosphorylation in both DNA replication and microtubule motor-related proteins, which cannot be accounted for by the initial disruption and which can explain in part the observed phenotype. Thus, the kinase is nevertheless expected to play a regulatory role in the seconds following activation. Furthermore, our observation that, as previously reported, microgametocytes lacking SRPK1 completely fail to exflagellate (Figure 5d; see also ref. (Tewari et al., 2010)), indicates that this kinase is required to further complete gametogenesis.

## Discussion

*Plasmodium* gametocytes must detect the change in environment from host to vector and propagate that signal quickly and efficiently to initiate the necessary gametogenesis processes. Because this process is vital to the parasite’s successful transmission, determining the underlying signalling events is a primary step towards a fundamental understanding of transmission. Our results reveal a prominent phosphorylation response within 18 s after gametocyte activation and that this signal is widespread by 60 s. In particular, despite having observed a large number of proteins undergoing active phosphoregulation during the response, we found a large fraction of the activity to occur in functionally related systems pertaining to DNA replication and mitosis, two processes known to be rapidly induced in the male gametocyte upon activation.

**Figure 5:**
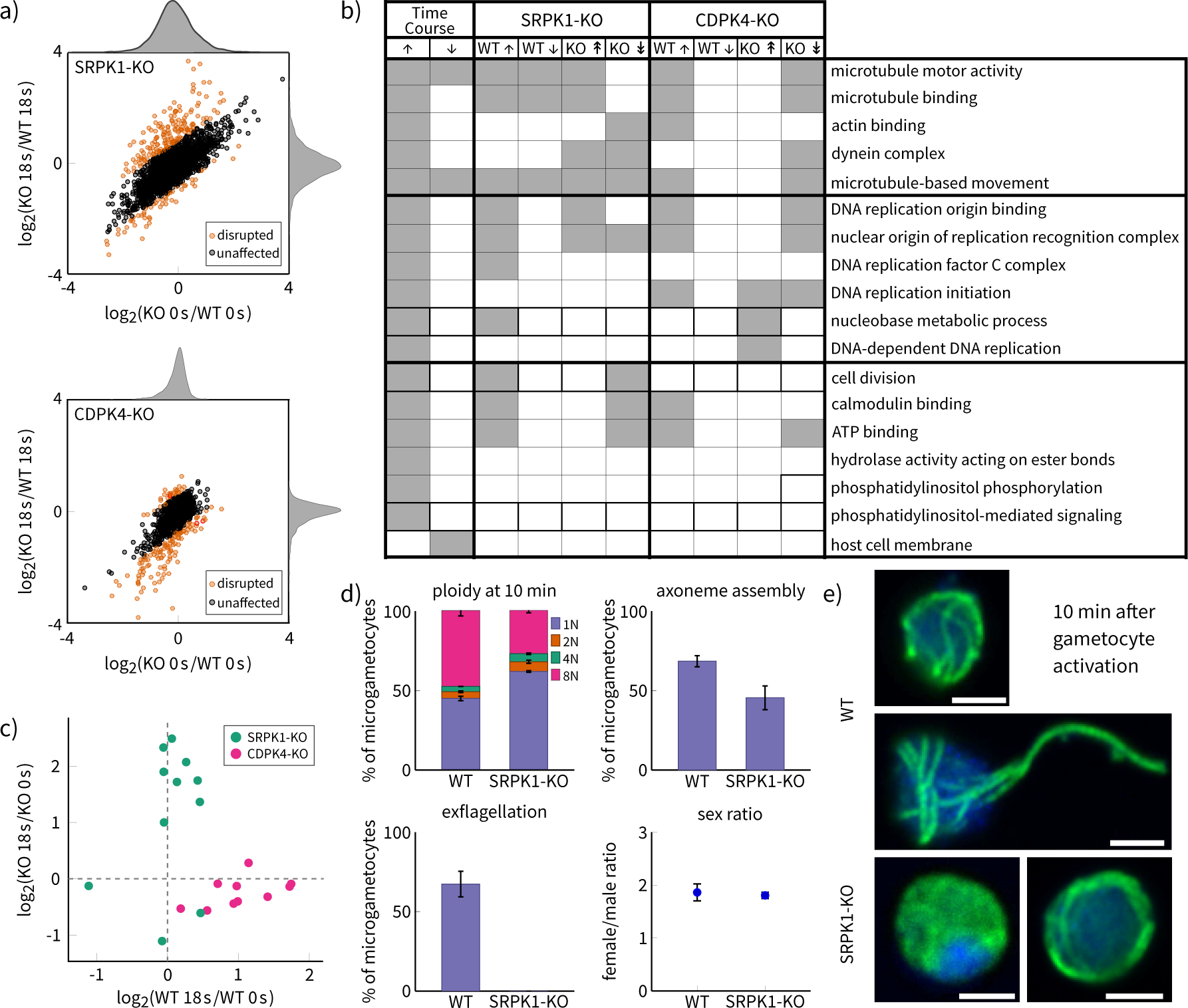
Individual deletions of the protein kinases CDPK4 and SRPK1 produce distinct effects on the phosphorylation response during early gametogenesis. a) Loss of CDPK4 or SRPK1 causes disruption versus wild type both before (0 s) and after (18 s) activation of the gametocytes. The degree of disruption in both states is greater in SRPK1-KO, as indicated by the distributions of fold-changes (marginal density plots). The outliers are peptides whose phosphoregulation was affected by the deletion following gametocyte activation. CDPK4-KO appears to have caused mainly loss of phosphorylation (negative disruption) while SRPK1-KO caused mainly gain of phosphorylation (positive disruption). b) A comparison of GO-term enrichment results for the time-course experiment and the two knock-out experiments (WT: up/down-phosphoregulation; KO: positive/negative disruption). Gray squares indicate significant enrichment. WT samples broadly replicate the time course experiment, given technical variation and the differences in experimental design. The KO samples show different effects on the same molecular systems. c) Disrupted phosphopeptides from motor proteins in the two KO experiments. Without SRPK1, several motor proteins become up-regulated during gametogenesis, while lack of CDPK4 results in the loss of phosphorylation events. d) The effect of knocking out SRPK1 on gametogenesis (two replicates; error bars represent standard error). Mutant parasites show reduced DNA replication and axoneme assembly, while exflagellation is completely blocked. The sex ratio was unaffected. e) Confocal microscopy images of WT (top, middle) and SRPK1-KO (bottom) parasites 10 min after activation. By this time, WT is initiating (top) or undergoing (middle) exflagellation, while some SRPK1-KO have exhibited no DNA replication or axoneme formation (bottom-left), or replication and axoneme formation but no exflagellation (bottom-right). Gametocytes were immunofluorescence-labelled for mouse anti-α-tubulin antibodies with anti-mouse Alexa488 as a secondary antibody (green). DNA was counterstained with DAPI (blue). Scale bars show 2 µm.

We observed phosphoregulation on several protein kinases and phosphatases, many of which are dispensable for the asexual erythrocytic cell cycle, but which were implicated through gene knockout studies in regulating gametocyte activation specifically, including CDPK1 and CDPK4, MAPK2, SRPK1, CnA, PPM1 and PPKL (Billker et al., 2004; Tewari et al., 2005; Rangarajan et al., 2005; Tewari et al., 2010; Sebastian et al., 2012; Guttery et al., 2014; Kern et al., 2014; Philip and Waters, 2015). In two cases, CDPK1 and CDPK4, we identified regulation of conserved phosphosites within their kinase domains, which may indicate regulation of their enzymatic activity during this process. For other kinases that we observed undergoing phosphoregulation, we cannot yet directly relate phosphorylation state to kinase activity levels. However, because some of these show significant functional association with other proteins undergoing phosphoregulation, we can infer that these kinases are likely to play important roles. In this manner, we identified ARK2, CRK5, NEK1, and the orphan kinase PBANKA_062210 as putative members of the gametocyte-activation signalling pathway. While no functional data are available on the orphan kinase, the first three candidates all belong to more conserved protein kinase subfamilies implicated in regulating the eukaryotic cell cycle. These have evidence for expression in asexual stages, and attempts to disrupt these genes have been consistently unsuccessful in *P. berghei* (Tewari et al., 2010; Solyakov et al., 2011). Roles for these protein kinases are therefore most likely not limited to gametogenesis, but may extend to regulating the *Plasmodium* cell cycle more generally. Stage-specific or inducible mutants in these genes have the potential to reveal deeper insights into the mechanisms of *Plasmodium* cell-cycle regulation.

Interestingly, another cdc2-related protein kinase, CRK4, is required for both replication and mitosis in *P. falciparum* schizonts and has been implicated in phosphorylation events on the pre-replicative complex. CRK4 is also expressed in *P. falciparum* gametocytes and while it is required for *Plasmodium* transmission to the mosquito, it is not essential for exflagellation (Ganter et al., 2017). Consistent with this, in *P. berghei* gametocytes we have observed phosphopeptides belonging to CRK1, 3, 4 and 5, but only the latter shows significant evidence of marked changes in phosphorylation levels, suggesting CRK5 is the likely S-phase promoting kinase in the activated microgametocyte. However, this does not rule out other potential roles for this kinase in gametogenesis. In *P. falciparum*, CRK5 is required for nuclear division during schizogeny (Dorin-Semblat et al., 2013), supporting the notion that CRK5 is central to parasite reproduction.

A comparison of the distinct effects of deleting CDPK4 and SRPK1 highlights the potential for complex modes of regulation in this system. Deletion of CDPK4 leads to a notable loss of phophorylation on many sites. CDPK4 thus appears to be directly or indirectly responsible for the phosphorylation of many proteins within the first 18 s, including those in the replisome and the microtubule cytoskeletal systems. Since CDPK4 is known to be activated in the microgametocyte soon after the XA signal is detected, this places this kinase upstream in the pathway with a “master regulator” role in the propagation of the signal, as previously suggested (Billker et al., 2004). SRPK1, on the other hand, plays a more indirect role, in which its presence is responsible for the repression of phosphorylation on proteins in the same systems. Taken together with our observation of reduced DNA replication and axoneme formation in the SRPK1 knock-out mutants, this implicates SRPK1 in gametocyte cell-cycle regulation rather than in its expected role of regulating SR-protein-mediated RNA processing. Because SRPK1 is itself negatively disrupted upon deletion of CDPK4, we hypothesise that SRPK1’s activity depends on activation of CDPK4 and that it, in turn, either negatively regulates another kinase or positively regulates a phosphatase. In at least one case, they appear to cooperate in a feedback-like manner. Considered in the light of the parasite’s reduced kinase repertoire (76 kinases in *P. berghei*) and the rapidity and efficiency of the gametocyte activation process, this provides an intriguing glimpse at the potentially complex signalling interactions underlying gametogenesis.

Cell-cycle regulation in malaria parasites is poorly understood and current evidence suggests that some of the canonical cell cycle check points are not present. During asexual schizogony in the blood stages, for example, nuclei behave as autonomous units that undergo repeated rounds of replication and mitosis in an asynchronous manner, while being retained within the same cellular envelope (Read et al., 1993; Arnot et al., 2011). During sexual development, the release of microgametes requires DNA synthesis (Janse et al., 1986), but on the other hand a compound that interferes with spindle formation did not prevent DNA replication to proceed through all three rounds (Billker et al., 2002), suggesting in the male nucleus multiple rounds of replication and mitosis may progress in parallel without depending on each other. We were therefore intrigued to see that protein modules involved in origin of replication recognition and cytoskeletal reorganisation were simultaneously targeted for phosphorylation during the first few seconds of gametocyte activation. These data indicate that some mitotic processes are already initiated contemporaneously with DNA synthesis in *Plasmodium* gametocytes and raise important questions regarding the temporal control of the cell cycle in these organisms.

## Materials & Methods

### Parasite maintenance and preparation

All animal experiments were conducted under a license from the UK Home Office in accordance with national and European animal welfare guidelines or with the authorisation number (GE/82/15) according to the guidelines and regulations issued by the Swiss Federal Veterinary Office. *P. berghei* strain ANKA (Vincke et al., 1966) derived clone 2.34 (Billker et al., 2004), CDPK4-KO (Fang et al., 2017), and SRPK1-KO (Tewari et al., 2010) were maintained in CD1 outbred mice obtained from Harlan or Charles River. For gametocyte production, mice were treated with phenyl hydrazine three days before infection. One day after infection asexually replicating parasites were eliminated by the addition of sulfadiazine (20 mg/L) in the drinking water. Parasites were harvested at day four after infection in suspended animation and separated from uninfected erythrocytes. Activation was induced and parasites were snap frozen in liquid nitrogen at 6, 12, 18 and 60 s after activation. For each time point (including unactivated “0 s”) and parasite line, four and two independent biological replicates were produced for the time-course experiment and the KO experiments, respectively. Cell samples were lysed and the protein content was purified and digested using the FASP method (Wiśniewski et al., 2009). See the Supplementary Methods for more details.

## Quantitative protein mass spectrometry

### Time course experiment

Phosphopeptide enrichment was performed on TiO_2_ tips (Thermo). Two technical replicates of each sample were analysed on an LTQ Orbitrap Velos coupled with an Ultimate 3000 RSLCnano System (both from Thermo Fisher). Raw spectra from the time-course experiment were analysed using MaxQuant (version 1.5.2.8) (Cox and Mann, 2008), treating technical replicates as fractions. Peptides were searched against the *P. berghei* annotated protein database retrieved from PlasmoDB (http://www.plasmodb.org; version 13.0) and the *M. musculus* protein sequence database retrieved from Uniprot (http://www.uniprot.org). Only phosphosites with a localisation probability of 0.75 or greater were retained for further analysis. For full protocol details and analysis parameters, see the Supplementary Methods.

Time courses for the change in phosphorylation level for each phosphosite were calculated using the composite intensity scores estimated by MaxQuant from all peptide evidence for that site. Following stringent quality control (see Supplementary Methods), three time courses were then generated from these values: the “full” time course (0–60 s) and two “truncated” timecourses for sites not detected in unactivated parasites (6–60 s) or at the end of the experiment (0–18 s). For each site, log_2_-transformed ratios were calculated from its intensity at each time point against its intensity at the first time point (*e.g.* log_2_ 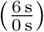). Only complete time courses, with no missing data, were retained for further analysis.

In order to determine which sites show significant evidence of change in phosphorylation state, we exploited the fact that enrichment of phosphopeptides is not 100% specific and that the time-frame in question is too short to see significant variation in protein abundances. Time courses were constructed for non-phosphorylated peptides in the same manner as described above. The distributions of non-phosphopeptide log_2_-ratios were used as null distributions against which the significance of the phosphosite log_2_-ratios could be tested. Sites having at least one time point with a *p*-value less than the critical value of 0.05 were determined to have undergone significant phosphoregulation during the time course. Significant sites were clustered according to their time courses using a normal mixture-modeling based method. See the Supplementary Methods for more details.

### Knock-out experiments

The CDPK4KO and the SRPK1KO experiments consisted of 8 samples each: wild-type and knockout samples measured at two time points, unactivated (0 s) and activated (18 s after activation), in two biological replicates. Cell samples were lysed and proteins were isolated via MTBE precipitation (Matyash et al., 2008). The proteins were then digested with trypsin and labelled with TMT 10plex. The labelled peptide mixture was fractionated and collected every 30 s and between 2.5–50 min on a 96-well plate by rows and then concatenated into 12 fractions by columns.

Enrichment of phosphopeptides was performed using IMAC with PHOS-Select Iron Affinity Gel (Sigma) then TiO_2_ tips (Thermo Fisher) sequentially. The enriched samples were subjected to LC-MS/MS analysis on an Orbitrap Fusion Tribrid mass spectrometer coupled with an Ultimate 3000 RSLCnano system configured as above. The phosphopeptides enriched via IMAC and TiO_2_ were analysed separately. Raw data were processed in Proteome Discoverer 2.1 (PD2.1; Thermo Fisher) using both SequestHT and Mascot search engines against a combined protein database of *P. berghei* and mouse as above. For full protocol details and analysis parameters, see the Supplementary Methods.

log_2_-ratios were calculated for both biological replicates using the peptides’ scaled abundances for the following comparisons for each replicate: KO 18 s vs. KO 0 s, WT 18 s vs. WT 0 s, KO 18 s vs. WT 18 s, and KO 0 s vs. WT 0 s. A final, mean ratio was computed from all of the available evidence (see Supplementary Methods). For all ratios, a *p*-value was calculated using the empirical cumulative distribution function of the non-phosphopeptides, as described above.

In order to account for a general change in cellular state before gametocyte activation, we built a linear model to predict the disruption at 18 s from the disruption at 0 s. That is, we aimed to predict 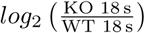 from 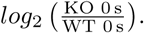. We reasoned that peptides which were poorly predicted by this model could be assumed to have been affected by the knock-out during the activation process. We then took the outliers to have been under direct influence of the deleted kinase (see Supplementary Methods for more details).

## Protein function and sex enrichment

Gene Ontology (GO) association files for *P. berghei* and *P. falciparum* (version 1.3.2015) were fetched from the Wellcome Trust Sanger Institute FTP server (ftp://ftp.sanger.ac.uk). The *P. falciparum* GO associations were used to supplement the *P. berghei* ones. For each *P. falciparum* gene with a one-to-one ortholog in *P. berghei*, the corresponding *P. falciparum* terms were merged with the existing terms associated with the *P. berghei* ortholog (if any). Groups of proteins were tested for enrichment of GO terms using GOATOOLS (version 0.5.9) (Tang et al., 2015), modified to perform the binomial test of significance (Mi et al., 2013). For the knock-out experiments, we only performed targeted enrichment analyses by testing only the “leaf” terms found enriched in the time-course experiment. For more details, see the Supplementary Methods.

Groups of proteins were further tested for enrichment for sex-specific proteins, using a previously published *P. falciparum* sex-partitioning data set by orthology (Tao et al., 2014). Fisher’s exact test was used to determine significant enrichment for female-specific, male-specific or shared proteins.

*p*-values for all enrichment analyses were corrected for FDR and significance was tested at a critical value of 0.05.

## Network analysis

Protein interaction networks for *P. berghei* and *P. falciparum* were downloaded from the STRING database (version 10; http://www.string-db.org) (Szklarczyk et al., 2015). The *P. berghei* network was extended to include additional nodes and edges from the *P. falciparum* network by orthology. New *P. berghei* nodes were added if they have a one-to-one ortholog in the *P. falciparum* proteome; edges from the *P. falciparum* STRING network were added to the *P. berghei* network only in the case that the edge was not already present; in the case that the STRING score of an edge between two orthologous pairs of proteins was different between the two networks, the original score from the *P. berghei* network was retained. Network statistics were computed using the NetworkX library for Python (version 1.8.1) (Hagberg et al., 2008).

Significant association on the network between protein kinases or phosphatases and the phosphosite time course clusters was determined through an enrichment analyses for each enzyme that was found to be phosphoregulated during the time course or that has previously been implicated in gametogenesis (see Figure 4). We compared the median of all of a kinase’s edge scores with the median of its edge scores shared with the proteins carrying phosphosites in a given time course cluster. We tested the hypothesis that the median edge score for the subset of proteins with sites in the cluster was higher than that of all the edges via the Mann-Whitney test. *p*-values were corrected for multiple testing by adjusting for FDR.

## Site conservation

Site conservation was measured by fitting nucleotide alignments of orthologous genes from seven *Plasmodium* species with site-specific codon substitution models. An estimate of the ratio of the number of non-synonymous substitutions per site to the number of synonymous substitutions per site (dN/dS) during species divergence was used as a measurement of conservation; dN/dS values less than one indicate that the site has predominantly been under purifying selection, with lower values indicating stronger selective constraint and conservation.

A nucleotide multiple-sequence alignment was produced for each genes in each orthologous group using PRANK (Löytynoja and Goldman, 2005; 2008). Regions with gaps were automatically removed using TrimAL (Capella-Gutiérrez et al., 2009). In order to estimate a species tree for the conservation analysis, a composite alignment was built by concatenating the 3150 alignments that contained orthologs from all seven species. The tree was estimated using PhyML (Guindon et al., 2010). Site dN/dS estimates were computed using the site models implemented in the “codeml” program of the PAML package (version 4.8a) (Yang, 2007). The analyses were automated using the PAML interface implemented in the Bio.Phylo module of Biopython (Talevich et al., 2012). Subsequent analyses of site conservation included only sites under purifying selection (dN/dS < 1). A regression model of dN/dS scores was built using beta regression and the residuals of this model were used to identify sites under stronger-than-expected conservation. For more details, see the Supplementary Methods.

## Motor protein phosphorylation hot-spots

We identified and grouped proteins belonging to the kinesin, dynein heavy chain, dynein light intermediate chain and dynein intermediate chain families according to their annotations in the proteome databases for *P. berghei*, *P. falciparum, T. gondii*, *M. musculus*, *Homo sapiens*, *Rattus norvegicus* and *Saccharomyces cerevisiae*. In addition, we considered a group of *Plasmodium* proteins for which no known function has been described (denoted as “conserved *Plasmodium* protein, unknown function”), but which have been assigned the “motor activity” GO term (GO:0003774) in *P. falciparum* at PlasmoDB, some of which we could assign by orthology to the kinesin family. Multiple sequence alignments were constructed for each family and phosphosite hot-spots were estimated via kernel density estimation on previously-published phosphosite counts at each aligned column. For more details, see the Supplementary Methods.

## Acknowledgements

Work at the Wellcome Trust Sanger Institute was funded by a core grant from the Wellcome Trust (WT098051). This work was supported by the Swiss National Science Foundation grant BSSGI0_155852 to MB. MB is an INSERM investigator. The mass spectrometry proteomics data have been deposited to the ProteomeXchange Consortium via the PRIDE (Vizcaíno et al., 2016) partner repository with the dataset identifier PXD006266. The authors declare no conflicts of interest.

## Author Contributions

OB, PB and JC supervised the work. OB, PB, JC, MB and BI designed the experiment. MB prepared the parasite cultures and performed the SRPK1-KO phenotyping. MB, BI and LY prepared the samples for LC-MS/MS analysis. LY performed the LC-MS/MS analysis. BI performed the data analyses and prepared the figures. All authors contributed to the manuscript text.

## Supplementary Methods

### Parasite maintenance and preparation

All animal experiments were conducted under a license from the UK Home Office in accordance with national and European animal welfare guidelines or with the authorisation number (GE/82/15) according to the guidelines and regulations issued by the Swiss Federal Veterinary Office. *P. berghei* strain ANKA (Vincke et al., 1966) derived clone 2.34 (Billker et al., 2004), CDPK4-KO (Fang et al., 2017), and SRPK1-KO (Tewari et al., 2010) were maintained in CD1 outbred mice obtained from Harlan. Mice were specific pathogen-free and subjected to regular pathogen monitoring by sentinel screening. They were housed in individually ventilated cages furnished with a cardboard fun tunnel and Nestlet. Mice were maintained at 21(2)°C under a 12 h light/dark cycle and given commercially prepared autoclaved dry rodent diet and water *ad libitum*. Mice were used for experimentation at 6 to 12 weeks of age.

For gametocyte production, mice were treated with phenyl hydrazine three days before infection. One day after infection asexually replicating parasites were eliminated by the addition of sulfadiazine (20 mg L^−1^) in the drinking water. Parasites were harvested at day four after infection in suspended animation (SA RPMI1640 medium containing25 mm HEPES, 5% FCS, 4 mm sodium bicarbonate, pH 7.2) and separated from uninfected erythrocytes on a Histodenz cushion made up from 48% of a Histodenz stock (27.6% w/v Histodenz -Sigma- in 5.0 mm Tris-HCl [pH 7.2], 3.0 mm KCl, 0.3 mm EDTA) and 52% SA with a final pH of 7.2. Purified gametocytes were washed twice in SA without FCS and resuspended in 400 µL of SA without FCS. Activation was induced by adding 400 µL of exflagellation medium (RPMI 1640 containing 25 mm HEPES, 4 mm sodium bicarbonate, 200 µm xanthurenic acid, pH 8). Parasites were snap frozen in liquid nitrogen at 6, 12, 18 and 60 s after activation. For the initial time point, the exflagellation medium was replaced by SA. For each time point and parasite line, four and two independent biological replicates were produced for the time-course experiment and the KO experiments, respectively.

A lysis buffer was prepared consisting of 4% SDS, 50 mm NaCl, 100 mm Tris buffer (pH 7.4), 5 mm EDTA, 40 mm TCEP, and Halt™ Protease & Phosphatase Inhibitor Cocktail (Thermo) (2 x). Samples were split in half (approximately 400 µL each) and 500 µL lysis buffer was added to each half. The samples were vortexed and then heated at 95°C for 10 min. DNA was sheared via pulses of sonication for 20 s (1 s on, 1 s off) at 40% power. The samples were centrifuged for 30 min at 14 000 rpm and the supernatant was collected. The samples were then incubated with 80 mm iodoacetamide for 1 h at room temperature. The protein digest used the FASP method (Wiśniewski et al., 2009) with Amicon Ultra-15 30K filter units (UFC903024), where Trypsin Gold (Thermo) was added at a 1:50 ratio and the digest was incubated at 37°C overnight. The resulting peptides were desalted using Sep-Pak C18 Plus Light cartridges then dried in a SpeedVac and stored at −20°C.

## Quantitative protein mass spectrometry

### Time course experiment

**Phosphopeptide enrichment** Phosphopeptide enrichment was performed on TiO_2_ tips (Thermo), following the manufacturer’s instructions. Each sample used 500 µg peptides. Phosphopeptides were eluted from the tip by 1.5% NH_4_OH followed by 5% pyrrolidine. Both eluates were pooled, acidified and then desalted on Graphite Spin Columns (Thermo) as instructed by the manufacturer’s protocol. Each sample was split into two technical replicates, dried in a SpeedVac, and stored at −20°C.

**Label-free LC-MS/MS** The dried phosphopeptides were resuspended in 80 µL of 0.5% FA / 100% H_2_O before LC-MS/MS analysis on an LTQ Orbitrap Velos coupled with an Ultimate 3000 RSLCnano System (both from Thermo Fisher). The peptides were first loaded and desalted on a PepMap C18 trap column (100 µm id x 20 mm, 5 µm) then separated on a PepMap C18 analytical column (75 µm id x 500 mm, 2 µm) (both from Thermo Fisher) over a 240 min linear gradient of 4 to 30% CH_3_CN/0.1% formic acid with the total cycle time at 280 min. The Orbitrap mass spectrometer was operated in the standard “top 15” data-dependent acquisition mode while the preview mode was disabled. The MS full scan was set at m/z 380 to 1600 with the resolution at 30 000 at m/z 400 and a lock mass at m/z 445.120 025. The AGC was set at 1 × 10^6^ with a maximum injection time at 200 ms. The 15 most abundant multiply-charged precursor ions, with minimal signals above 3000 counts, were dynamically selected for CID fragmentation (MS/MS) in the ion trap, which had the AGC set at 5000 with the maximum injection time at 100 ms. The dynamic exclusion duration time was set for 60 s with ±10 ppm exclusion mass width. The isolation width was 2.0 Da and the normalised collision energy was 35%. LC-MS/MS analyses of technical replicates were repeated sequentially.

**Spectral analysis and peptide quantification** Raw spectra from the time-course experiment were analysed using MaxQuant (version 1.5.2.8) (Cox and Mann, 2008). Technical replicates were treated as fractions. The minimum peptide length was set at seven and protein and site identification false discovery rates (FDR) were set at 0.01. Carbamidomethyl was set as a fixed modification. Peptides were searched against the *P. berghei* annotated protein database retrieved from PlasmoDB (http://www.plasmodb.org; version 13.0) and the *M. musculus* protein sequence database retrieved from Uniprot (http://www.uniprot.org). Peptides were quantified using the iBAQ (intensity-based absolute quantification) method (Schwanhäusser et al., 2011), as implemented in MaxQuant, using a logarithmic fit and spectra-matching between runs. Acetylation, deamidation, oxidation and phosphorylation were included as modifications in the protein quantification. Only phosphosites with a localisation probability of 0.75 or greater were retained for further analysis.

**Data processing** Time courses for the change in phosphorylation level for each phosphosite were calculated using the composite intensity scores estimated by MaxQuant from all peptide evidence for that site. For each time point (0, 6, 12, 18 and 60 s), these phosphosite intensities were quantile normalised between the four biological replicates Bolstad et al. (2003). For each site at each time point, the mean and standard error of the intensities were calculated if intensities were measured in at least two replicates, otherwise it was considered not to have been detected. The standard error of the site’s intensities at a given time point was compared to the distribution of standard errors of all sites’ intensities for that time point. If a site’s standard error of intensities at any time point was greater than the distribution’s upper quartile plus 1.5 times the interquartile range (IQR) (*i.e.* the upper “whisker” on a standard box-and-whisker plot), that site was discarded as unreliable.

Three time courses were then generated from these values: the “full” time course (0 to 60 s), for sites that were detected at all five time points; a “late” truncated time course (6 to 60 s), for sites that were not detected in unactivated parasites; and an “early” truncated time course (0 to 18 s), for sites that were not detected at the 60 s time point. For each site, ratios were calculated from its intensity at each time point against its intensity at the first time point and were logarithmically transformed (base 2) (*e.g.* 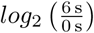). Only complete time courses, with no missing data, were retained for further analysis.

In order to determine which sites show significant evidence of change in phosphorylation state, we exploited the fact that enrichment of phosphopeptides is not 100% specific and that the time-frame in question is too short to see significant variation in protein abundances. Time courses were constructed for non-phosphorylated peptides in the same manner as described above. All phosphosite ratios and nonphosphopeptide ratios were then corrected for the median non-phosphopeptide ratio for the corresponding time point. The distributions of non-phosphopeptide log_2_ ratios were used as null distributions against which the significance of the phosphosite log_2_ ratios could be tested. Two-tailed *p*-values were computed from an empirical cumulative distribution function of the non-phosphopeptide ratios. Sites showing at least one time point with a *p*-value less than the critical value of 0.05 were determined to have undergone significant phosphoregulation during the time course.

Significant sites were clustered according to their time courses using a normal mixture-modeling based method, using the “mclust” library (version 5.2) in R (Fraley and Raftery, 2002; Fraley et al., 2012). The library chooses the number of clusters and the specific model to fit (spherical with equal volumes or spherical with unequal volumes) via an automated method which optimises the Bayesian Information Criterion (BIC).

### Knock-out experiments

**Phosphopeptide enrichment** The CDPK4KO and the SRPK1KO experiments consisted of 8 samples each: wild-type and knock-out samples measured at two time points, unactivated (0 s) and activated (18 s after activation), in two biological replicates.

200 µL of 2 x lysis buffer was added to the 200 µL parasite suspension, and then processed as described above. 300 µg proteins were taken and the volumes were equalised with 100 mm TEAB followed by alkylation with IAA. Proteins were precipitated by MTBE (Matyash et al., 2008) and then digested with 8 µg trypsin in 150 µL of 100 mm TEAB at 37°C for 2 h. A further 4 µg trypsin was added and digested for another 5 h. 100 µg peptides was taken from each replicate for TMT 10plex labelling, then mixed and dried in a SpeedVac. The labelled peptide mixture was fractionated at pH 10 on an XBridge BEH C18 column (4.6 mm id x 250 mm, 130 Å, 3.5 µm) (Waters) at a flow rate of 500 µL min^−1^ with a linear gradient from 5 to 35% ACN/NH_3_ in 30 min and total cycle time of 60 min. Fractions were collected every 30 s and between 2.5 to 50 min on a 96-well plate by rows and then concatenated into 12 fractions by columns and dried in a SpeedVac.

Enrichment of phosphopeptides was performed using IMAC with PHOS-Select Iron Affinity Gel (Sigma) then TiO_2_ tips (Thermo Fisher) sequentially. All procedures followed the manufacturer’s instruction with some modification. The peptides were redissolved in 50% ACN/0.1% TFA then added to 100 µL of pre-washed PHOS-Select Iron Affinity Gel and left binding at room temperature with end-to-end rotation for 30 min. The beads were washed three times with 250 mm acetic acid/50% ACN and once with H_2_O. Phosphopeptides were eluted twice with 100 µL of 1.5% NH_3_/25% ACN then dried in a SpeedVac. The flow-through and the first wash of IMAC beads were collected and dried in a SpeedVac, and then the phosphopeptides were enriched using TiO_2_ tips as described above.

**TMT-labelled LC-MS/MS analysis** The enriched phosphopeptides were redissolved in 0.5% FA before LC-MS/MS analysis on an Orbitrap Fusion Tribrid mass spectrometer coupled with an Ultimate 3000 RSLCnano system configured as above. The peptides were separated with a linear gradient of 4 to 36% ACN/0.1% FA in 120 min and total 153 min per cycle. The Orbitrap Fusion was operated using the “Top Speed” method with 2 s cycle time. The MS full scan was in the Orbitrap with the following settings: scan range at m/z 380 to 1500 with a lock mass at 445.120 025, resolution at 120 000 at m/z 200, and AGC at 4 × 10^5^ with a maximum injection time at 50 ms. The multiply-charged precursor ions (2+ to 6+) at most intense, with a minimal signal above 10 000 counts, were dynamically selected for high energy collision-induced dissociation (HCD) (MS/MS) and detected in the Orbitrap with a resolution at 30 000 at m/z 200. The isolation width was 1.2 Da in quadrupole, and the collision energy was set at 40%. The dynamic exclusion duration time was set for 60 s with ±10 ppm exclusion mass width, the AGC was set at 1 × 10^5^ with the maximum injection time at 105 ms.

The phosphopeptides enriched via IMAC and TiO_2_ were analysed separately.

**Spectral analysis and peptide quantification** Raw data were processed in Proteome Discoverer 2.1 (PD2.1; Thermo Fisher) using both SequestHT and Mascot search engines against a combined protein database of *P. berghei* and mouse as above. Trypsin maximum missed cleavage sites were set to 2. The dynamic modifications set in both Mascot and SequestHT were Acetyl (N-term), Deamidated (NQ), Phospho (STY) and Oxidation (M), while in SequestHT Camabidomethyl (C) was set as a fixed modification. The remaining settings were the same in both: precursor mass tolerance at 20 ppm, fragment at 0.5 Da, and TMT6plex as fixed modification. All files acquired from same set of TMT10plex experiments, including from both IMAC and TiO_2_ purified samples, were set as fractions. The search result was validated by Percolate where the q-value was set at 0.01. The PSMs were identified, quantified and grouped to the peptides. Both PSMs and peptides were filtered with 1% FDR, then further group to proteins with 1% FDR where only peptides at high confidence were selected. Both unique and razor peptides were used for protein quantification, and protein and peptide abundance values were calculated as summed PSM quan values (S/N values of reporter ions). The abundances were normalised on Total Peptide Amount, and then scaled with On Channels Average. The co-isolation threshold was set at 50% to reduce the isolation interference. The phosphorylation sites were localised by phosphoRS as implemented in PD2.1 with site probability set at 75.

**Data processing** For each knock-out experiment, peptides were filtered to include those which were unambiguously matched to a single *P. berghei* or *M. musculus* protein with a FDR less than 0.01. Furthermore, only peptides that were quantified in all TMT channels were retained. log_2_ ratios were calculated for both biological replicates using the peptides’ scaled abundances provided by Proteome Discoverer for the following comparisons: KO 18 s vs. KO 0 s, WT 18 s vs. WT 0 s, KO 18 s vs. WT 18 s, and KO 0 s vs. WT 0 s. For each comparison in each biological replicate, the ratios were corrected by subtracting the median value. Finally, for each comparison, a mean ratio was calculated by first computing the mean ratio within each replicate for each of the peptide’s PSMs (*i.e.* those PSMs with phosphorylation at the same site(s) but otherwise differing in other modifications), followed by computing the mean of the resulting value between the two biological replicates. For all ratios, a *p*-value was calculated using the empirical cumulative distribution function of the non-phosphopeptides, as described above.

In order to account for a general change in cellular state before gametocyte activation, we built a linear model to predict the disruption at 18 s from the disruption at 0 s. That is, we aimed to predict 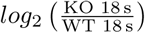 from 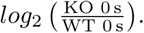 We reasoned that peptides which were poorly predicted by this model could be assumed to have been affected by the knock-out during the activation process. To quantify this, we Studentised the residuals and assigned *p*-values from the t-distribution. Peptides that were found to have both residual *p*-values and 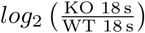 ratio *p*-values less than 0.05 were taken to be significantly affected by the knockout. We also performed a further partitioning of the affected peptides to isolate those which suffered an apparent loss of phospho-regulation, by taking those which were: a) observed to be significantly up-regulated in the WT sample or in an up-regulation cluster in the time-course experiment; and b) either significantly down-regulated or not significantly regulated in the KO sample.

## Determination of gametocyte ploidy by FACS analysis

Ploidy of gametocytes was determined by FACS measurement of fluorescence intensity of cells stained with Vybrant dye cycle violet (life Technologies). Parasites were treated for two days with sulfadizine and gametocytes were purified and resuspended in 100 µL of SA. Activation was induced by adding 100 µL of modified exflagellation medium (RPMI 1640 containing 25 mm HEPES, 4 mm sodium bicarbonate, 5% FCS, 200 µm xanthurenic acid, pH 7.8). To rapidly block gametogenesis, 800 µL of ice-cold PBS was added and cells were stained for 30 min at 4°C with Vybrant dye cycle violet. Cells were analysed with a Beckman Coulter Gallios 4. Per sample, fluorescence intensity of >50 000 cells was determined with the Kaluza analysis software.

## Immunofluorescence labelling and microscopy

Gametocyte immunofluorescence assays were performed as previously described (Volkmann et al., 2012). Purified cells were fixed with 4% paraformaldehyde and 0.05% glutaraldehyde in PBS for one hour, permeabilised with 0.1% Triton X-100/PBS for 10 min and blocked with 2% BSA/PBS for 2 h. Mouse anti-α-tubulin clone DM1A (Sigma-Aldrich) primary antibodies were diluted 1:1000 in blocking solution. Anti-mouse Alexa488 was used as secondary antibody together with DAPI (all from Life technologies), all diluted 1:1000 in blocking solution. Confocal images were acquired with a LSM700 scanning confocal microscope (Zeiss).

## Orthology

Protein orthology relationships were predicted using OMA (standalone version 0.99z.2) (Roth et al., 2008). The orthology predictions were performed on the proteomes of seven *Plasmodium* species: *P. berghei* ANKA, *P. chabaudi chabaudi*, *P. falciparum* 3D7, *P. knowlesi* H, *P. reichenowi* CDC, *P. vivax* Sal1, and *P. yoelii yoelii* 17XNL. Annotated protein sequences were fetched from PlasmoDB (version 13.0; http://www.plasmodb.org). OMA was run with a minimum alignment score of 181, an alignment length tolerance of 0.41, a stable-pair distance tolerance of 1.81, a verified-pair distance tolerance of 1.53 and a minimum sequence length of 50 residues.

## Protein function enrichment

Gene Ontology (GO) association files for *P. berghei* and *P. falciparum* (version 1.3.2015) were fetched from the Wellcome Trust Sanger Institute FTP server (ftp://ftp.sanger.ac.uk). The *P. falciparum* GO associations were used to supplement the *P. berghei* ones. For each *P. falciparum* gene with a one-to-one ortholog in *P. berghei*, the corresponding *P. falciparum* terms were merged with the existing terms associated with the *P. berghei* ortholog (if any).

Groups of proteins were tested for enrichment of GO terms via a binomial test (Mi et al., 2013). Query protein groups for the time-course experiment included the list of all proteins containing significant sites and the lists of proteins containing significant sites in each of the phosphosite clusters. For the knockout experiments, the query groups included proteins that showed significant up- or down-regulation in the wild-type sample, those whose post-activation state was significantly disrupted in the knock-out versus the wild-type, and those which we determined to be putative down-stream targets of the deleted kinase. The “background” set of proteins, against which enrichment was tested, consisted of all of the proteins detected in the time-course mass-spectrometry experiments, including those detected via non-phosphorylated peptides. GO enrichment analyses were performed using GOATOOLS (version 0.5.9) (Tang et al., 2015), modified to perform the binomial test, on the “go-basic” ontology file provided at http://purl.obolibrary.org/obo/go/go-basic.obo. For the knock-out experiments, we only performed targeted enrichment analyses by testing only the “leaf” terms found enriched in the time-course experiment. *p*-values for all enrichment analyses were corrected for FDR and significance was tested at a critical value of 0.05.

## Protein Domain Prediction

The presence of Pfam domains in *P. berghei* proteins was predicted using the “pfamscan” script (upload date 2013-10-15) available on the Wellcome Trust Sanger Institute FTP server (ftp://ftp.sanger.ac.uk/pub/databases/Pfam/Tools). The script performed sequence searches using version 3.1b1 of HMMR Eddy (2011) and version 27.0 of the Pfam database (Finn et al., 2016).

## Site Disorder

Site disorder estimates were calculated using DISOPRED (version 3.16) (Jones and Ward, 2003). DISOPRED performed PSI-BLAST searches using the BLAST executable “blastpgp” (version 2.2.26) on the “nr” database with the default parameters specified in the script provided by the DISOPRED software package.

## Site Conservation

Site conservation was measured by fitting nucleotide alignments of orthologous genes with site-specific codon substitution models. An estimate of the ratio of the number of non-synonymous substitutions per site to the number of synonymous substitutions per site (dN/dS) during species divergence was used as a measurement of conservation; dN/dS values less than one indicate that the site has predominantly been under purifying selection, with lower values indicating stronger selective constraint and conservation.

A nucleotide multiple-sequence alignment was produced for the genes in each orthologous group using PRANK (version .140110), aligning by codons (option “-codon”) and leaving sites inferred as insertions unaligned (option “-F”) (Löytynoja and Goldman, 2005; 2008). Regions with gaps were automatically removed using TrimAL (Capella-Gutiérrez et al., 2009). In order to estimate a species tree for the conservation analysis, a composite alignment was built by concatenating the 3150 alignments that contained orthologs from all seven species. The tree was estimated using PhyML with the GTR nucleotide substitution model and the better of the NNI and SPR tree topology search methods (option “-s BEST”) (Guindon et al., 2010).

Site *dN/dS* estimates were computed using the site models implemented in the “codeml” program of the PAML package (version 4.8a) (Yang, 2007). The analyses were automated using the PAML interface implemented in the Bio.Phylo module of Biopython (Talevich et al., 2012). In order to distinguish between positive and purifying selection, each alignment was fitted with a pair of models, termed M8 and M8a (Swanson et al., 2003; Wong et al., 2004). These models are nested, permitting the better model to be chosen via a likelihood-ratio test. Model M8a assigns sites to classes either with an estimated 0 < *dN/dS* < 1 drawn from the *β*-distribution (purifying selection) or with *dN/dS* = 1 (neutral evolution). Model M8 is similar, albeit with the latter category estimated with *dN/dS* > 1 (positive selection). The equilibrium codon frequencies in the models were estimated from the average nucleotide frequencies at the three codon positions. The parameter was estimated from an initial value of 4.0, the parameter was fixed at infinity, and the parameter *ρ* was fixed at 0. The likelihood-ratio tests were performed with one degree of freedom using twice the log-likelihood difference of the two models on the 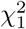 distribution (Wong et al., 2004); a significant test would indicate that the M8 model, featuring positively selected sites, better fits the alignment than the null model, M8a. *p*-values were adjusted for FDR and a critical value of 0.05 was used to determine significance. dN/dS estimates for specific sites on a given protein (*e.g.* significant phosphosites) were taken from the Naive Empirical Bayes (M8a) or Bayes Empirical Bayes (M8) site-rate predictions for the chosen model, as produced by the software.

Subsequent analyses of site conservation included only sites under purifying selection (*dN/dS* < 1). Because we focus only on sites with 0 < *dN/dS* < 1, the data is best represented by the beta distribution. A regression model of *dN/dS* scores was built using beta regression, as implemented in the “betareg” library for R (version 3.0-5) (Cribari-Neto and Zeileis, 2010). Two terms are modelled: the mean and a precision parameter, *ϕ*, which accounts for heteroskedasticity. After model reduction, the coefficients in the model of the mean were: DISOPRED score; median *dN/dS* score for the twenty sites preceding and twenty sites following the site, not including the three nearest positions on either side; the log_10_-transformed median expression level of the RNA encoding the protein (Otto et al., 2014); whether or not the site is significantly regulated; and an interaction between the region median *dN/dS* and the median expression level terms. The mean model was fit using the logit link function. The model of the precision parameter included the following coefficients: DISOPRED score, log_10_-transformed median RNA expression level, and whether or not the site is significantly regulated. All terms were significant at a level of *p* < 0.001.

## Motor Protein Phosphorylation Hot-Spots

We identified and grouped proteins belonging to the kinesin, dynein heavy chain, dynein light intermediate chain and dynein intermediate chain families according to their annotations in the proteome databases for *P. berghei*, *P. falciparum, T. gondii*, *M. musculus*, *Homo sapiens*, *Rattus norvegicus* and *Saccharomyces cerevisiae*. In addition, we considered a group of *Plasmodium* proteins for which no known function has been described (denoted as “conserved *Plasmodium* protein, unknown function”), but which have been assigned the “motor activity” GO term (GO:0003774) in *P. falciparum* at PlasmoDB. We performed orthology predictions as described above for all of these protein families. Several of the unannotated *Plasmodium* proteins were found to be members of a hierarchical orthology group which contained annotated kinesin-like proteins, including the human proteins KIF15 and KIF20B: PBANKA_123650, PBANKA_093930, PBANKA_092720, PBANKA_121680, PBANKA_112530, and PBANKA_124420. We thus treated these proteins and their *P. falciparum* orthologs as belonging to the kinesin family. Previously published human, mouse and rat phosphosite information was retrieved from Phosphosite Plus (Hornbeck et al., 2015) (date of data set: 2016 Jan. 29). For yeast, phosphosite data was retrieved from Phosphogrid (Sadowski et al., 2013) (downloaded 2014 Mar. 14). Phosphosite data was manually curated for *T. gondii* (Treeck et al., 2011; 2014), *P. falciparum* (Solyakov et al., 2011; Treeck et al., 2011; Lasonder et al., 2012; Pease et al., 2013; Collins et al., 2014; Lasonder et al., 2015) and *P. berghei* (Brochet et al., 2014). The *P. berghei* was further supplemented with the phosphosites observed in our time course experiment.

A multiple sequence alignment was computed for each family using MAFFT (version 7.205) (Katoh and Standley, 2013), using the “E-INS-i” methodology, which is “suitable for sequences containing large unalignable regions” (options “--ep 0 --genafpair --maxiterate 1000”). Alignment quality was calculated using Jalview (version 2.9.0b2) (Waterhouse et al., 2009), normalised by the maximum quality, and a running mean was calculated using a window of length 15. Hot-spots were estimated by performing a kernel density estimation on the phosphosite counts at each column in the alignment, using the Epanechnikov kernel and using the Sheather & Jones (“SJ”) algorithm for automatic bandwidth selection (Sheather and Jones, 1991), as implemented in R.

## Network Visualisation

Network visualisations were produced using Ti*k*Z and LATEX based on a custom modification of the Walshaw 2000 spring-electrical layout algorithm (Walshaw, 2000; 2003; Tantau, 2013), using a spring constant of 0.1, a convergence tolerance of 0.001, an electric force order of 4, an electric charge of 0.085, and a cooling factor of 1.0. Phosphosite nodes were added to protein nodes as a sub-layout using the same algorithm with a spring constant of 1.5 and an electric force order of 1 (fewer than nine phosphosites) or 0 (nine or more phosphosites).

## Supplementary Figures

**Figure 1:**
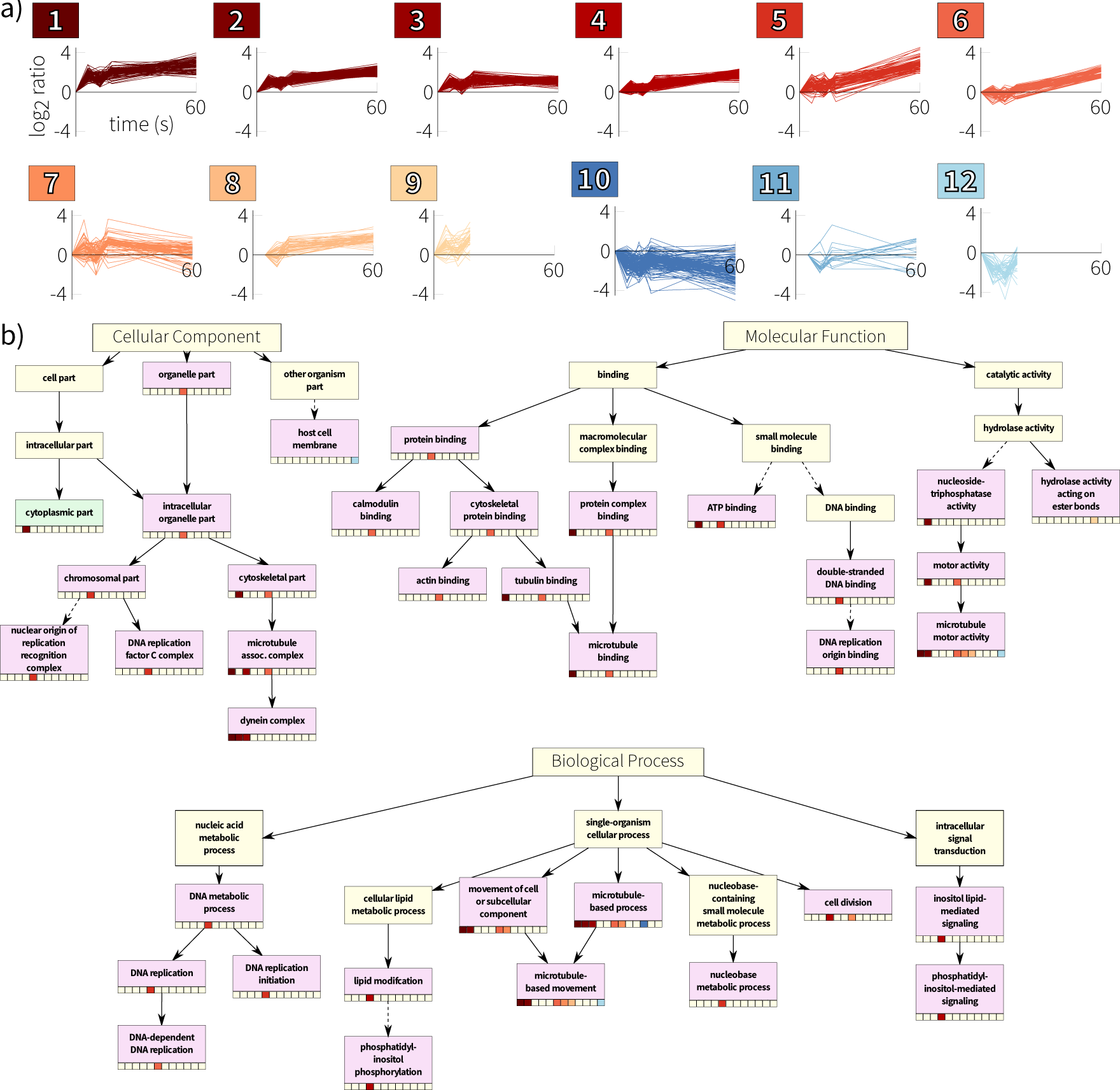
a) Gaussian mixture-model-based clustering of P. berghei phosphorylation time courses. b) Enriched (pink) and depleted (green) Gene Ontology terms for the P. berghei time-course clusters, including enrichment for more general, parent terms (higher in the graph of terms). For each such term, the clusters enriched/depleted for the term are indicated in the cells beneath the term. The arrangement of the cluster cells, from left to right, matches the numbering in panel (a). Note that some intermediate terms have been removed for clarity; relationships which include removed terms are indicated by dashed lines.

**Figure 2:**
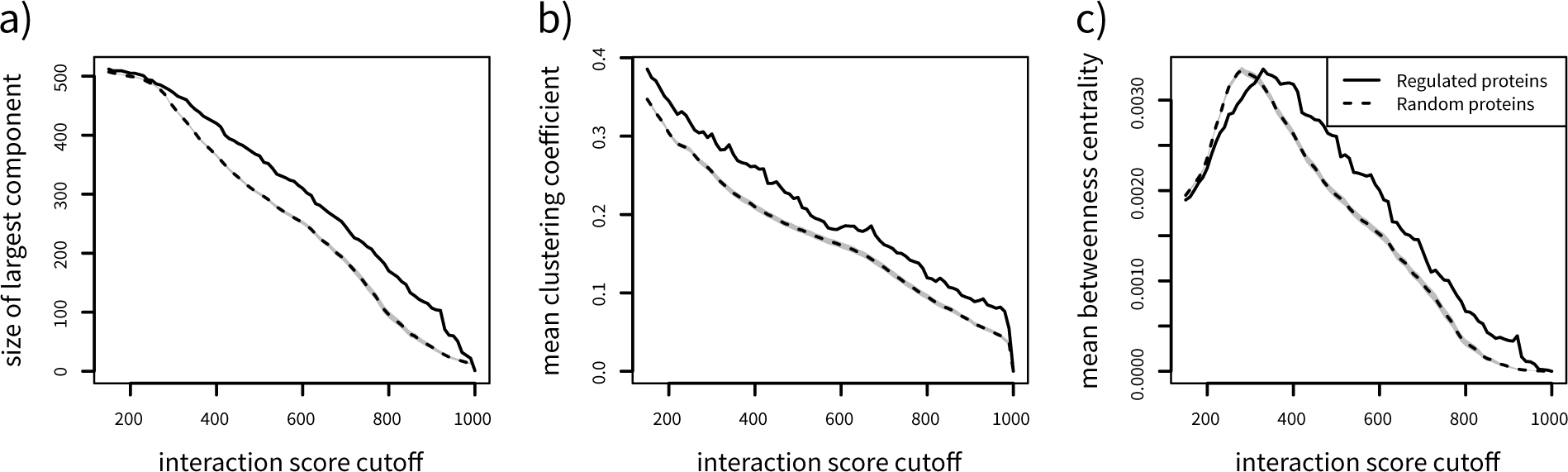
The subset of the P. berghei protein-protein association network that consists only of proteins that were phosphoregulated in the time course experiment shows a) a larger connected component, a) higher median clustering coefficient, and c) higher median betweenness centrality across the full range of interaction score cutoff values than random subsets of the same size. The shaded grey area indicates the 95% confidence interval of the measurements for the random sub-networks.

**Figure 3:**
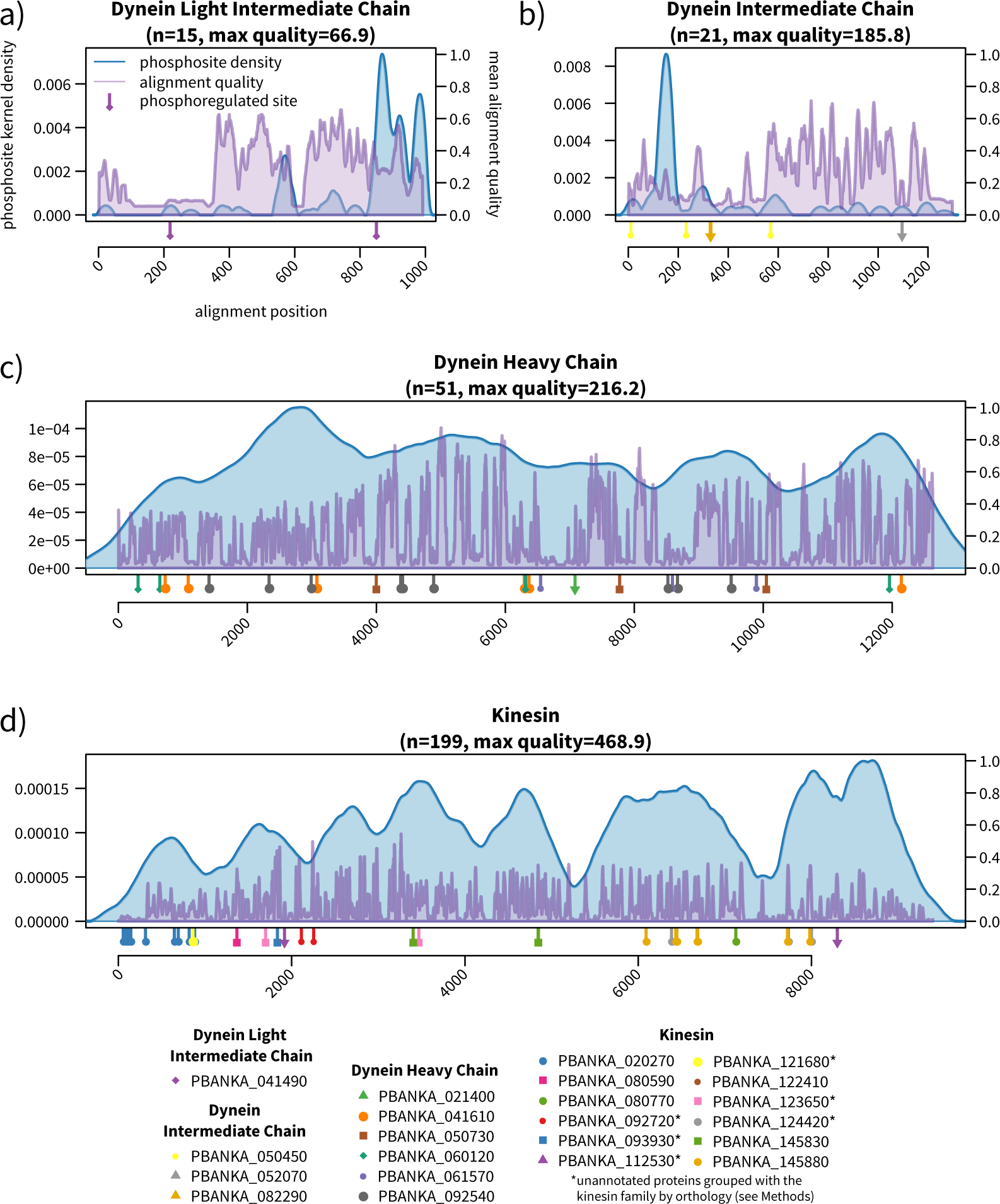
A comparison of regulated phosphosite positions with phosphorylation patterns across motor protein families. Blue traces show the kernel density estimation of phosphosite counts across members of the protein family from P. berghei, P. falciparum, T. gondii, H. sapiens, M. musculus, R. norvegicus, and S. cerevisiae. Violet traces show a running mean of the normalised alignment quality. The pins beneath the plots show the locations of P. berghei sites that were observed to be under phosphoregulation during gametocyte activation. Dynein light intermediate (a) and intermediate chains (b) have regulated phosphosites at or near phosphorylation hot-spots. Dynein heavy chain (c) and kinesin (d) show broad phosphorylation across their entire lengths, with regulated sites being relatively evenly.

